# Age-related pathological impairments in induced neurons derived from patients with idiopathic Parkinson’s disease

**DOI:** 10.1101/2021.04.23.441070

**Authors:** Janelle Drouin-Ouellet, Karolina Pircs, Emilie M. Legault, Marcella Birtele, Fredrik Nilsson, Shelby Shrigley, Maria Pereira, Petter Storm, Yogita Sharma, Romina Vuono, Thomas B. Stoker, Johan Jakobsson, Roger A. Barker, Malin Parmar

## Abstract

Understanding the pathophysiology of Parkinson’s disease has been hampered by the lack of models that recapitulate all the critical factors underlying its development. Here, we generated functional induced dopaminergic neurons (iDANs) that were directly reprogrammed from adult human dermal fibroblasts of patients with idiopathic Parkinson’s disease to investigate disease-relevant pathology. We show that iDANs derived from Parkinson’s disease patients exhibit lower basal chaperone-mediated autophagy as compared to iDANs of healthy donors. Furthermore, stress-induced autophagy resulted in an accumulation of macroautophagic structures in induced neurons (iNs) derived from Parkinson’s disease patients, independently of the specific neuronal subtype but dependent on the age of the donor. Finally, we found that these impairments in patient-derived iNs lead to an accumulation of phosphorylated alpha-synuclein, a hallmark of Parkinson’s disease pathology. Taken together, our results demonstrate that direct neural reprogramming provides a patient-specific model to study aged neuronal features relevant to idiopathic Parkinson’s disease.

## Introduction

Parkinson’s disease is a neurodegenerative disorder that has as part of its core pathology the loss of midbrain dopaminergic (DA) neurons and the aggregation of the misfolded protein alpha-synuclein (αsyn). How the disease arises and develops is currently unknown and no cure exists. There is an urgent need for better treatments and disease modifying therapies, but their development is hampered by a poor understanding of the pathogenesis of Parkinson’s disease and lack of appropriate model systems, in particular ones which capture age – the single biggest risk factor for developing this condition – as well as the heterogeneity of idiopathic Parkinson’s disease.

The idiopathic nature of most Parkinson’s disease cases (>90%), coupled to the late age of onset, complicates experimental studies of the pathophysiology and it is challenging to design and interpret models of idiopathic Parkinson’s disease. For example, most animal models depend on toxin-induced mitochondrial damage or overexpression of αsyn at high levels (i.e. levels that exceed that seen in the brain of patients dying with idiopathic Parkinson’s disease). Patient-derived induced pluripotent stem cells (iPSCs) are frequently used to study cellular features of Parkinson’s disease^1–3^ but fail to capture the age and epigenetic signatures of the patient.^4–7^ In order to better recapitulate these disease-relevant features, we and others have developed protocols for the generation of subtype-specific induced neurons (iNs) that can be directly generated from human fetal fibroblasts using specific combinations of transcription factors and fate determinants.^8–12^ This type of direct neural conversion offers several advantages. In particular, directly reprogrammed cells have been shown to retain many important aspects of the donor-derived starting fibroblasts, including age-related changes in the epigenetic clock, the transcriptome and microRNAs, the reactive oxygen species level, extent of DNA damage and telomere length, as well as metabolic profile, mitochondrial defects and protein degradation defects.^6,13–18^

In this study, we identified a combination of reprogramming factors that resulted in the generation of subtype-specific and functional induced DA neurons (iDANs) when converting dermal fibroblasts from aged individuals. Subsequently, we used this protocol to convert iDANs from idiopathic Parkinson’s disease patient-derived fibroblasts as well as age- and sex-matched controls. When analyzing the patient-derived neurons, we found that stress-induced chaperone-mediated autophagy (CMA) and macroautophagy impairments could be detected in the idiopathic Parkinson’s disease iNs but not in control iNs nor in parental fibroblasts of the patients. This type of pathology has previously mainly been captured in genetic Parkinson’s disease variants using iPSC-based models.^19–23^ This study thus reports Parkinson’s disease-associated phenotypes in directly converted neurons from patient fibroblasts and provides a new model to study idiopathic forms of Parkinson’s disease.

## Results

### Generation of functional iDANs from dermal fibroblasts of adult donors

In order to establish a model of Parkinson’s disease using iNs with characteristics of DA neurons, we screened 10 different reprogramming factors (*Ascl1, Lmx1a, Lmx1b, FoxA2, Otx2, Nurr1, Smarca1, CNPY1, EN1, PAX8*) that were selected based on their; (i) role during normal DA neurogenesis^34^; (ii) expression in human fetal ventral midbrain^35^; (iii) value in predicting functional DA differentiation from stem cells^36^, and/or (iv) role on midbrain-specific chromatin modeling.^37^ All factors were expressed in combination with the knockdown of the REST according to our published protocol for high efficiency reprograming of adult fibroblasts.^24^ Three of the screened combinations gave rise to a significant proportion of TH expressing neurons: shREST + Ascl1 + Lmx1a/b + FoxA2 + Otx2 (2.2 % ± 2.0), shREST + Ascl1 + Lmx1a/b + FoxA2 + Otx2 + Smarca1 (7.5 % ± 3.6), and shREST + Ascl1 + Lmx1a/b + FoxA2 + Otx2 + Nurr1 (**Fig. 1a; Supplementary Fig. 1a**). The best TH-positive cell yield was obtained with the last combination: shREST + Ascl1 + Lmx1a/b + FoxA2 + Otx2 + Nurr1. This combination gave rise to up to 70.3 % ± 0.3 of cells expressing the neuronal marker TAU, of which 16.1 % ± 2.01 also expressed TH (**Supplementary Fig. 1a**), with an average TAU purity of 9.1% ±3.3 and TH purity of 2.6% ±1.7 (**Fig. 1b,c**), and showed robust upregulation of DA genes as measured by qRT-PCR (**Supplementary Fig. 1b**).

**Figure 1.**
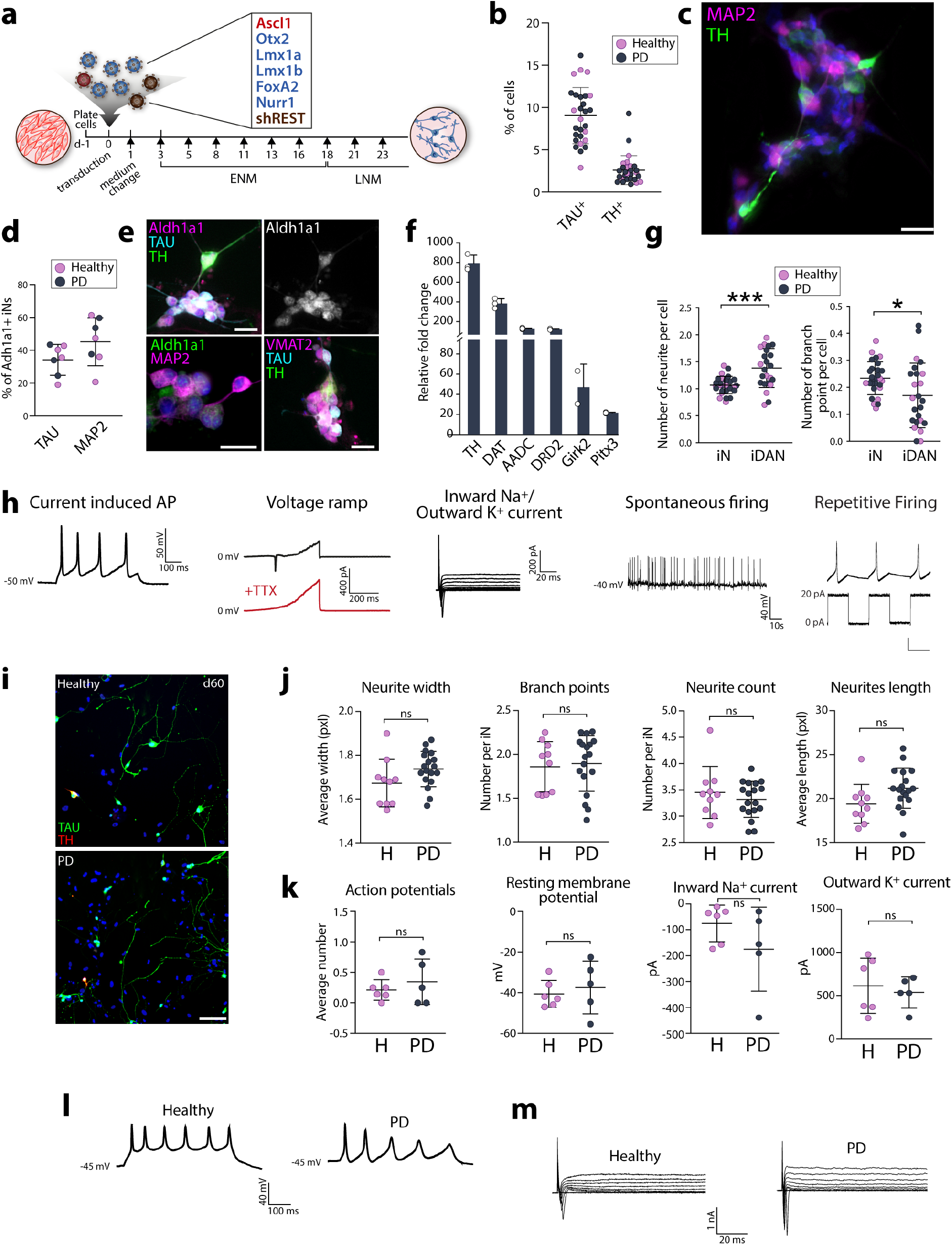
Generation of iDANs from Parkinson’s disease and heathy donor lines. **a**, Reprogramming iDANs from adult fibroblasts. **b**, Quantification of TAU-positive and TH-positive cells (mean average of 2,575 TAU-positive and 32 TH-positive cells assessed per line, n=28 lines). **c**, MAP2-positive and TH-positive iDANs. Cells are counterstained with DAPI (in blue). Scale bars = 25μm. **d**, Quantification of ALDH1A1 and TAU or MAP2 double positive cells (mean average of 1,652 TAU-positive and 1,258 MAP2-positive cells assessed per well from 4 biological replicates per lines, n=7 lines (lines #4, #8, #9, #10, #26, #27 and #28). **e**, TAU-positive, MAP2-positive and TH-positive iNs and iDANs expressing ALDH1A1 and VMAT2. Cells are counterstained with DAPI (in blue). Scale bars = 25μm. **f**, Gene expression quantification of DA genes relative to parental fibroblast levels (from 2 to 3 biological replicates (white circles) from line #2). **g**, Quantification of the neurite profile in TAU-positive and TH-negative (iNs) vs. TAU and TH double positive cells (iDANs) from healthy and Parkinson’s disease lines (mean average of 2,575 TAU-positive and 32 TH-positive cells assessed per line, n=28 lines). Two-tailed unpaired t-test with Welch’s correction: ***P=0.0004, df=30.82; *P=0.0245, df=32.94. **h**, Patch clamp recordings of iDANs from line #2 (at day 65). **i**, Double TAU-positive and TH-positive H-iDANs and PD-iDANs at day 60. Scale bar = 100μm. **j**, Quantification of the neurite profile in TAU-positive H-iNs and PD-iNs. (experiment has been repeated independently 3 times, mean average of 2,142 TAU-positive cells assessed per line, n=10 healthy and n=18 Parkinson’s disease lines). **k**, Quantification of voltage-clamp recordings of evoked action potentials (*n* = 8-10 neurons per lines, *n* = 5-6 lines per group), resting membrane potential of H-iNs and PD-iNs. (*n* = 4-9 neurons per lines, *n* = 5-6 lines per group), inward and outward currents (*n* = 4-9 neurons per lines, *n* = 5-6 lines per group). Lines #1, #2, #4, #5, #6, #8, #13, #16, #17, #24, #28 were used for patch clamp experiments. **l**, Voltage-clamp recordings of repetitive evoked action potentials. **m**, Representative traces of membrane sodium- and potassium currents following voltage depolarization steps in H-iNs and PD-iNs. Abbreviations: APs: action potentials, ns: not significant, TTX: tetrodotoxin.

Further characterization of the iNs obtained using this reprogramming factor combination showed that 35.4% ± 9.1 of the TAU-positive and 45.2% ± 14.7 of the MAP2-positive cells also expressed ALDH1A1 (**Fig. 1d**), which is found in a subset of A9 DA neurons that are more vulnerable to loss in Parkinson’s disease,^38^ as well as VMAT2, a key DA neuronal marker (**Fig. 1e**). Gene expression profiling of 76 neuronal genes related to dopaminergic, glutamatergic and GABAergic neuronal subtypes confirmed an up-regulation of key genes related to DA patterning and identity (*FOXA1, OTX1, SHH, PITX3*), as well as DA synaptic function - including the receptors *DRD1* to *DRD5*, the DA transporter *DAT*, the enzymes *DDC, MAOA, ALDH1A1* and the A9-enriched DA marker *GIRK2* (**Fig. 1f; Supplementary Fig. 1c, d**). Thus the iDANs expressed the key markers of the AT-DAT^high^ subgroup of the DA sublineage as identified in Tiklová et al.,^39^ (expressing ALDH1A1 and TH, and localized mainly in the SNc), including the transgenes Lmx1a, Lmx1b, Foxa2, and Nurr1 (Nr4a2), and also DAT, VMAT2, AADC and Pitx3. In addition, given that a proportion of the iNs generated were TH-negative and given that Ascl1-based conversion protocols have been shown to generate excitatory and inhibitory neurons,^40^ we also looked for expression of genes associated with glutamatergic and GABAergic neurons. We found high expression of many of these genes, including vGlut1, GABAARb3, GAD, AMPAR2, Kainate R subunit 2 25 days after initiation of conversion (**Supplementary Fig. 1d**), suggesting a mixed culture of these neuronal subtypes. Finally, to get a better idea of the identity of the cells that are not iNs, we have performed a triple staining using MAP2 to identify iNs, as well as GFAP and Collagen 1 to identify potential glial cells and cells that remained fibroblasts. We observed that some MAP2-negative cells are expressing either Collagen 1 or GFAP alone, or together (**Supplementary Fig. 1e**).

Morphological assessment showed that TH-positive iNs express significantly more neurites compared to non-TH iNs but had significantly less branch points (**Fig. 1g**). Patch-clamp electrophysiological recordings 65 days post transduction confirmed that the reprogrammed cells had functionally matured (**Fig. 1h**). They were able to fire repetitive action potentials upon injection of current as well as exhibited inward sodium - outward potassium currents with depolarizing steps. When a continuous depolarizing voltage ramp was applied, inward currents was seen across the membrane and could be blocked by the neurotoxin tetrodotoxin (TTX), indicating an involvement of voltage-gated sodium channels in these currents. Without any injection of current or voltage, the cells displayed spontaneous firing and 43.8% of iNs also showed rebound action potentials and/or pacemaker like activity typical of mesencephalic DA neurons (**Fig. 1h**). Based on this, we refer to the cells as induced DA neurons, iDANs.

### Generation of functional iDANs from dermal fibroblasts of idiopathic Parkinson’s disease patients

Using this new iDAN reprogramming method, we next converted fibroblasts obtained from skin biopsies of 18 idiopathic Parkinson’s disease patients and 10 age- and sex-matched healthy donors (See **Table 1**). We found that the fibroblasts obtained from Parkinson’s disease patients reprogrammed at a similar efficiency to those obtained from healthy donors (**Fig. 1b**) and displayed a similar neuronal morphological profile (**Fig. 1i, j**). Moreover, when measuring their functional properties using patch clamp electrophysiological recordings, we confirmed that iNs derived from healthy donors (H-iNs) and from Parkinson’s disease patients (PD-iNs) displayed similar functionalities in terms of the number of current induced action potentials, resting membrane potential and the inward sodium - outward potassium currents (**Fig. 1k-m**).

**Table 1.** Demographics, clinical and genotype data of the study participants.

| Line ID | Group | Sex | Age at biopsy | MAPT haplotype | Age at onset | Disease duration (years) | *UPDRS motor decline | *MMSE score decline |
| --- | --- | --- | --- | --- | --- | --- | --- | --- |
| 1 | Healthy | M | 69 | H1/H1 |  |  |  |  |
| 2 | Healthy | F | 67 | H1/H1 |  |  |  |  |
| 3 | Healthy | M | 80 | H2/H2 |  |  |  |  |
| 4 | Healthy | F | 75 | H1/H1 |  |  |  |  |
| 5 | Healthy | M | 70 | H1/H2 |  |  |  |  |
| 6 | Healthy | F | 70 | H1/H2 |  |  |  |  |
| 7 | Healthy | M | 71 | H1/H1 |  |  |  |  |
| 8 | Healthy | F | 61 | H1/H2 |  |  |  |  |
| 9 | Healthy | F | 66 | H2/H2 |  |  |  |  |
| 10 | Healthy | F | 58 | H1/H1 |  |  |  |  |
|  | Ratio F:M/<br>Mean±SD | 6:4 | 68.7 ±<br>6.3 |  |  |  |  |  |
| 11 | PD | M | 56 | H1/H1 | 34 | 23 | -0.32 | 0 |
| 12 | PD | M | 60 | H1/H1 | 48 | 12 | -0.43 | 0 |
| 13 | PD | F | 77 | H2/H2 | 65 | 12 | 1.48 | 0 |
| 14 | PD | F | 67 | H1/H1 | 56 | 12 | 0.76 | -0.07 |
| 15 | PD | F | 59 | H1/H1 | 45 | 15 | -1.39 | 0 |
| 16 | PD | F | 80 | H1/H2 | 69 | 11 | 0.39 | -0.07 |
| 17 | PD | M | 80 | H2/H2 | 49 | 33 | 1.70 | 0 |
| 18 | PD | F | 87 | H1/H1 | 72 | 15 | 0.25 | -0.12 |
| 19 | PD | F | 77 | H1/H1 | 56 | 24 | -0.42 | -0.25 |
| 20 | PD | M | 75 | H1/H1 | 63 | 13 | -1.18 | 0 |
| 21 | PD | M | 77 | H1/H1 | 66 | 11 | -1.68 | 0 |
| 22 | PD | F | 71 | H1/H1 | 62 | 14 | 0 | -0.47 |
| 23 | PD | M | 72 | H1/H1 | 70 | 2 | 1.71 | -0.44 |
| 24 | PD | M | 81 | H1/H1 | 76 | 6 | 1.88 | 0.14 |
| 25 | PD | F | 44 | H1/H1 | 40 | 5 | -3.56 | 0.15 |
| 26 | PD | F | 79 | H1/H1 | NA | NA | NA | NA |
| 27 | PD | F | 68 | H1/H1 | 55 | 15 | 1.65 | 0 |
| 28 | PD | M | 57 | H1/H1 | 50 | 8 | NA | NA |
|  | Ratio F:M/<br>Mean±SD | 10:8 | 70.4 ±<br>11.2 |  | 57.4<br>±<br>11.9 | 12.5 ±<br>7.1 | 0.05 ±<br>1.5 | -0.07±<br>0.18 |

\* Average rate of decline per year over a minimum of 2 years.
Abbreviations: MMSE: Mini-Mental State Examination; NA: not available; UPDRS: Unified Parkinson's Disease Rating Scale

### PD-iDANs show altered chaperone-mediated autophagy

To assess the presence of age- and Parkinson’s disease-related pathological impairments in iDANs derived from idiopathic Parkinson’s disease patients, we focused on autophagy, a lysosomal degradation pathway that is important in cellular homeostasis and the efficiency of which decreases with age.^41^ We first looked for CMA alterations as this is one type of autophagy that has been suggested to be implicated in the pathophysiology of Parkinson’s disease.^42^ During CMA, HSC70 recognizes soluble, cytosolic proteins carrying a KFERQ-like motif and guides these proteins to the transmembrane LAMP2a receptor.^43^ Thereafter the protein cargo is translocated into the lysosomal lumen and as such, the level of LAMP2a determines the rate of CMA. ^44^ To induce autophagy, cells were cultured under starvation conditions, which promotes the recycling of non-essential proteins and organelles for reuse. ^45^ After validating that the starvation regimen had no impact on the number of neurons and induced changes in LAMP2a and HSC70 expression using WB (**Supplementary Fig. 2a-c**), we assessed CMA expression using a high content screening approach, which allowed us to analyze cytoplasmic puncta in parental fibroblasts, iNs (TAU-positive and TH-negative) and iDANs (TAU-positive and TH-positive) in a quantitative manner, and also to determine their subcellular location. When investigating this in parental fibroblasts and PD-iNs at baseline and in the context of starvation using an antibody specific to the “a” isoform of LAMP2, we did not observe a difference in LAMP2a-positive cytoplasmic puncta in parental fibroblasts between H-iNs and PD-iNs (**Supplementary Fig. 3a,b)** nor in the neurites of TAU-positive iNs upon starvation in both H-iNs and PD-iNs (**Supplementary Fig. 4a,b**). However, when looking specifically at iDANs, we observed a lower number of LAMP2a-positive cytoplasmic puncta in the neurites at baseline in the PD-iDANs compared to H-iDANs, suggesting a lower basal rate of CMA in PD-iDANs (**Fig. 2a,b**). Importantly, the decrease in LAMP2a-positive cytoplasmic puncta seen in TAU-positive neurites was only present in iDANs from the healthy group of donors, suggesting that PD-iDANs have an altered response to starvation, and that this alteration is specific to the DA subtype (**Fig. 2a,b**). We next looked at HSC70 expression, the main chaperone responsible for the degradation of αsyn via CMA.^46^ Both parental fibroblasts from healthy and Parkinson’s disease donors showed a decrease in HSC70 expression in response to starvation (**Supplementary Fig. 3c,d**). While the number of HSC70-positive puncta in the neurites of starved H-iDANs increased (154.0 % ± 117.2 of the nonstarved condition), starvation-induced autophagy led to a decrease of HSC70-positive puncta in PD-iDANs (55.9 % ± 28.1 of the non-starved condition) (**Fig. 2c**). Taken together, and in line with what has been previously reported in animal models of Parkinson’s disease,^47,48^ these results suggest that there is both an alteration in baseline CMA as well as stress-induced autophagy that is specific to idiopathic Parkinson’s disease derived iDANs.

**Fig 2.**
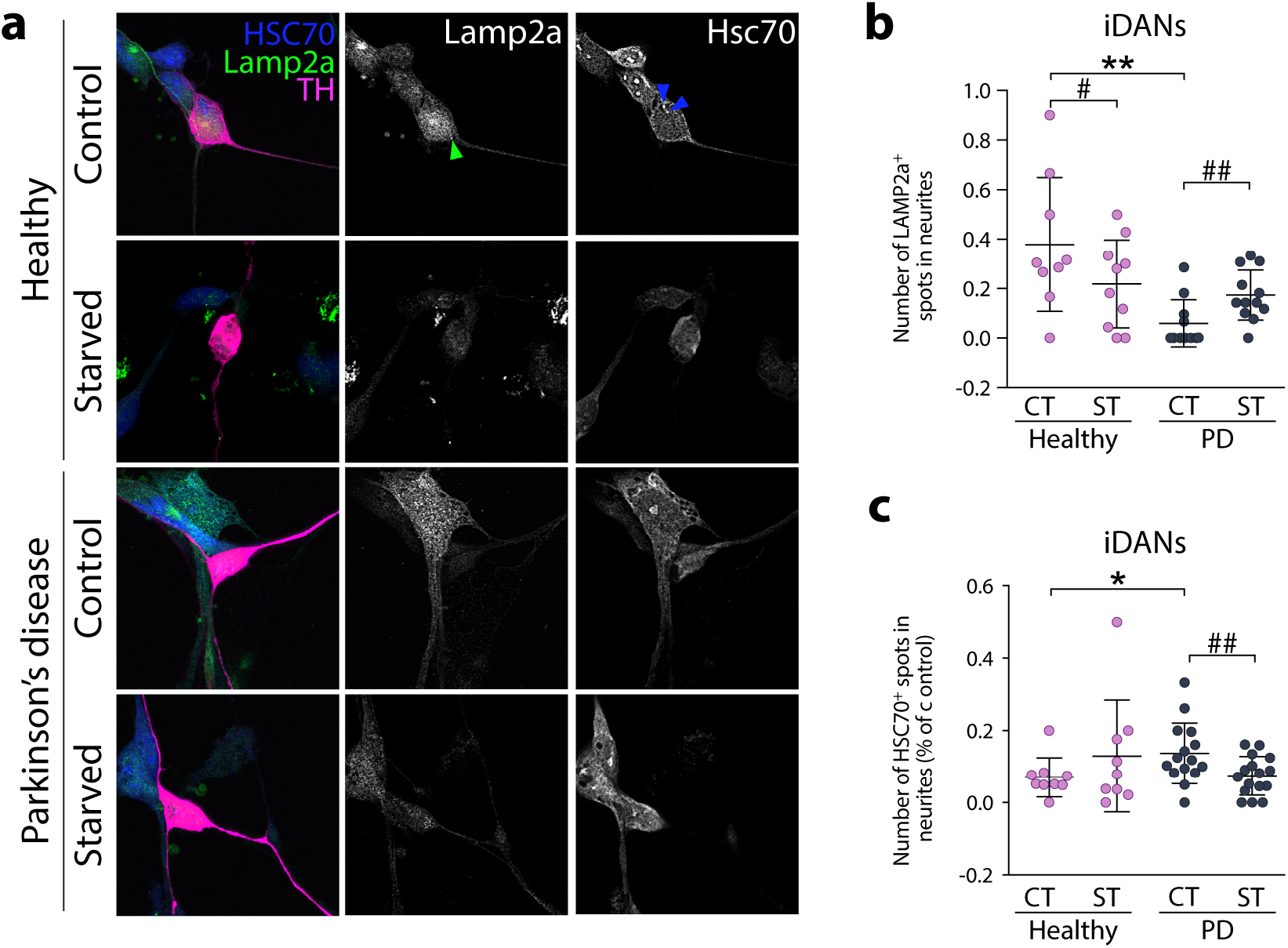
Chaperone-mediated autophagy impairment in PD-iDANs. **a**, LAMP2a-positive dot expression and spot detection analysis of LAMP2a-positive (green arrowhead) and HSC70-positive (blue arrowheads) puncta in TH-positive iDANs. **b**, Quantification of LAMP2a-positive puncta in the neurites of TH-positive iDANs (mean average of 14 TH-positive cells assessed per line, n=10 healthy and n=18 Parkinson’s disease lines). Kruskal-Wallis test, Dunn’s multiple comparisons test: *P=0.0067; H: Two-tailed paired t-test: ^#^P=0.0194, df=8; Parkinson’s disease: Wilcoxon matched pairs signed rank test: ^##^P=0.0098, rs=0.339. **c**, Quantification of HSC70-positive puncta in neurites of TH-positive iDANs (mean average of 95 TH-positive cells assessed per line, n=8-9 healthy and n=16 Parkinson’s disease lines). Mann-Whitney *U* test: *P=0.0128, *U*=26.5. Wilcoxon matched pairs signed rank test: ^##^P=0.0031, rs=0.395. Abbreviations: CT: control, H: healthy, PD: Parkinson’s disease, ST: starved.

### Altered macroautophagy response to stress-induced autophagy in iNs from Parkinson’s disease patients

CMA preferentially degrades specific proteins, rather than organelles and other macromolecules.^49,50^ However, while there is considerable cross talk between CMA and macroautophagy, starvation predominantly induces macroautophagy - a process involving the formation of double membraned autophagosomes which fuse with lysosomes, resulting in degradation of their contents. Given that we observed CMA alteration in PD-iDANs in response to starvation, we sought to further investigate whether there is an impairment in macroautophagy in PD-iNs. To validate the activation of macroautophagy upon starvation, we first looked at the cargo receptor p62, which decreases in the context of nutrient deprivation.^51^ In the parental dermal fibroblasts, our starvation regimen induced a decrease in p62-positive cytoplasmic puncta in both healthy and Parkinson’s disease donor derived lines (**Supplementary Fig. 5a, b**). This decrease was also observed in H-iNs (70.6 % ± 27.2 of the non-starved condition in the cell body and 77.1% ± 22.8 in neurites) (**Fig. 3a, b**). However, once converted to neurons, the majority of Parkinson’s disease lines failed to degrade p62 upon starvation, resulting in an accumulation of p62-positive puncta in PD-iNs as compared to H-iNs, which was observed in all TAU-positive iNs, regardless of the neuronal subtype or neuronal compartment (115.3 % ± 38.4 of the non-starved condition in the cell body and 108.4% ± 47.1 in neurites) (**Fig. 3a, b**). We then assessed more specifically LC3 to identify autophagic structures.^30^ We found that starvation significantly reduced the size of LC3-positive cytoplasmic puncta in the cell bodies of H-iNs (26.5 % ± 15.9 of the non-starved condition). However, LC3-positive cytoplasmic puncta in the cell bodies of PD-iNs were not significantly smaller after starvation (56.5 % ± 44.2 of the non-starved condition). When comparing the level of size reduction of LC3-positive cytoplasmic puncta in the cell bodies after starvation, PD-iNs failed to reduce dot size to the level that was seen in the H-iN group (**Fig. 3c, d**), whereas no effect of starvation was observed in the neurites of H-iNs and PD-iNs (116.3 % ± 53.6 of the non-starved condition for H-iNs, and 104.4 % ± 51.8 for PD-iNs). Furthermore, this difference in macroautophagy between H-iNs and PD-iNs in the cell bodies was a cell type specific feature as it was not seen in the parental fibroblasts (**Supplementary Fig. 5c, d**).

**Fig 3.**
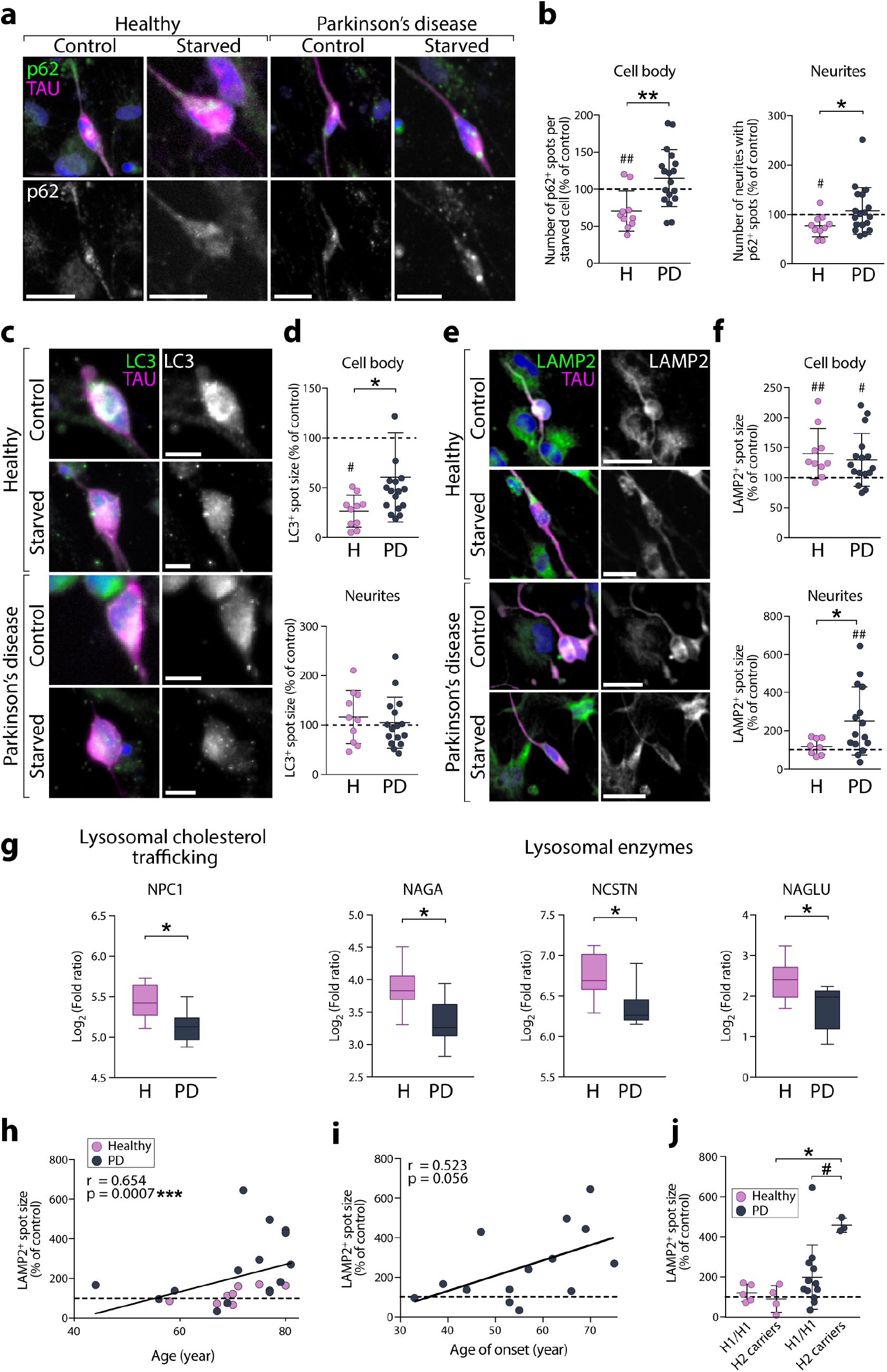
Accumulation of p62, LC3 and LAMP2 in PD-iNs upon starvation. **a**, p62-positive dot expression in TAU-positive iNs. Scale bar = 25μm. **b**, Quantification of p62-positive puncta in TAU-positive iNs (mean average of 577 TAU-positive cells assessed per line, n=10 healthy and n=18 Parkinson’s disease lines). Cell body: Two-tailed unpaired t-test: **P=0.0032, df=26, H: two-tailed paired t-test: ^##^P=0.0056, df=9; Neurite: Mann-Whitney *U* test: *P=0.0452, *U*=48, H: two-tailed paired t-test: ^#^P=0.0128, df=9. Data were normalized as % of control condition (not starved). **c**, LC3-positive dot expression in TAU-positive iNs. Scale bar = 10μm. **d**, Quantification of LC3-positive puncta in TAU-positive iNs (mean average of 479 TAU-positive cells assessed per line, n=10 healthy and n=18 Parkinson’s disease lines). Cell body: two-tailed Mann-Whitney *U* test: *P=0.0311, *U*=51, H: two-tailed paired t-test: ^##^P=0.0051, df=9. Data were normalized as % of control condition (not starved). **e**, LAMP2-positive dot expression in TAU-positive iNs. Scale bar = 25μm **f**, Quantification of LAMP2-positive puncta in TAU-positive iNs (mean average of 202 TAU-positive cells assessed per line, n=10 healthy and n=17 Parkinson’s disease lines). Cell body: H: two-tailed paired t-test: ^##^P=0.0078, df=8, Parkinson’s disease: ^#^P=0.0295, df=13. Neurites: Two-tailed unpaired t-test with Welch’s correction: *P=0.0136, *U*=24: *P=0.0125, df=18.23, Parkinson’s disease: twotailed paired t-test: ^##^P=0.0042, df=16.75. Data were normalized as % of control condition (not starved). **g**, Boxplots of log2 fold changes in expression of genes associated with lysosomal functions (adjusted P value < 0.09, n=10 healthy and n=10 Parkinson’s disease lines). **h**, accumulation of LAMP2-positive puncta upon stress-induced autophagy is associated with the age of the donor (n=23 lines). Spearman’s rank correlation: ***P=0.0007; 95% confidence interval: 0.3199 to 0.8437. **i**, association between accumulation of LAMP2-positive puncta upon stress-induced autophagy and the age of onset of Parkinson’s disease (n=15 Parkinson’s disease lines). Spearman’s rank correlation: *P=0.0431; 95% confidence interval: 0.01127 to 0.8263. **j**, More pronounced accumulation of LAMP2-positive puncta upon stress in MAPT H2 carrier Parkinson’s disease patients. Kruskal-Wallis test, Dunn’s multiple comparisons test: *P=0.0265; two-tailed Mann-Whitney *U* test: ^#^P=0.0250, *U*=3. Abbreviations: CT: control, H: healthy, PD: Parkinson’s disease, ST: starved.

Once autophagosomes have enclosed their autophagy substrates, they can fuse with endosomes or lysosomes to form amphisomes and autolysosomes. We thus used LAMP2 (detecting all three isoforms: LAMP2a, LAMP2b and LAMP2c) to visualize these structures. LAMP2-positive cytoplasmic puncta decreased upon starvation in the parental fibroblasts of Parkinson’s disease lines (**Supplementary Fig. 5e, f**). However, while an increase of the size of these structures upon starvation was similar in the cell bodies of both H- and PD-iNs (140.1 % ± 41.6 of the non-starved condition for H-iNs and 130.06 % ± 43.8 for PD-iNs), in the neurites, the size of LAMP2-positive puncta was unaffected by starvation in H-iNs (118.9 % ± 42.3 of the non-starved condition), whereas they were significantly bigger in PD-iNs (251.8 % ± 177.7 of the non-starved condition) (**Fig. 3e, f**). Unlike the altered CMA response (**Fig. 2a-c**), these phenotypes were present in all iNs and not just DA neurons (**Supplementary Fig. 6**).

To assess whether this altered autophagy response could be due to basal changes in the transcriptome of PD-iNs, we performed RNA-seq analysis on the iNs and the parental fibroblasts. This analysis confirmed a profound change in gene expression profile as fibroblasts were reprogrammed towards a neuronal transcriptome (**Supplementary Fig. 7**). Moreover, GSEA using KEGG pathways identified genes in the lysosome pathway (hsa014142) to be significantly enriched (adjusted p-value = 0.026) (**Supplementary Fig 8a**).When analyzing specifically the lysosomal genes, we found that the lysosomal cholesterol trafficking gene *NCP1* involved in the inherited metabolic disease Niemann-Pick, type C,^52^ as well as three other lysosomal enzymes (NAGA, NCSTN, NAGLU) were down-regulated in PD-iNs compared to H-iNs (**Fig. 3g**), supporting the data suggesting that there are alterations in lysosomal functions at baseline and is in line with observations that these inherited disorders can lead to parkinsonian states clinically.^53^ Importantly, when analyzing expression of these genes between the healthy controls and Parkinson’s disease patients in parental fibroblasts, they were not differentially expressed (**Supplementary Fig. 8b**).

We next sought to determine the impact of blocking the autophagic flux in starved cells using Bafilomycin A1, which inhibits the fusion of endosomes/lysosomes with autophagosomes/ amphisomes.^44^ Here again, we observed a different response in PD-iNs (108.9 % ± 28.8 of the starved condition), which accumulated significantly less LC3-positive cytoplasmic dots upon inhibition of autophagy in the cell body as compared to H-iNs (146.1 % ± 56.8 of the starved condition) (**Supplementary Fig. 8c-e**). This suggests there is an impairment in the early phases of autophagy. In support of this, differential gene expression analysis of autophagy-related genes (ATG) revealed that multiple genes involved in the early autophagy processes were significantly changed at baseline in the PD-iNs. These were genes involved in the autophagy initiation ATG1 kinase complex and regulators (*ATG13, CAMK2B*), in the ATG12 (*ATG7* and *FBXW7*) and ATG8 conjugation systems (*ATG7*), and the ATG2-ATG18 complex (*WDR45*) (**Supplementary Fig. 8f**). This was also accompanied by a down-regulation of *TM9SF1*, that is involved in autophagosome formation and which also interferes with starvation-induced autophagy when down-regulated in cells (**Supplementary Fig. 8d**).^54^ These genes were not differentially expressed in the parental fibroblasts (**Supplementary Fig. 8e**), consistent with the absence of autophagic alterations observed in the starting cells (**Supplementary Fig. 5a-d**).

### Age-related correlation in disease-associated impairments and accumulation of phosphorylated αsyn

Recent reports have shown that age-associated properties of the human donors are maintained in iNs but not in iPSC-derived neurons.^6,15–18^ We therefore assessed if the accumulation of lysosomal structures in H- and PD-iNs was associated with the age of the donor. In addition to a positive correlation between the age and the accumulation of lysosomes in neurites (**Fig. 3h**), we also found a trend towards a positive correlation of this accumulation with age of onset at diagnosis (**Fig. 3i**). This was more pronounced in lines derived from patients carrying the H2 haplotype of *MAPT*, which has previously been associated with a more rapid progression and cognitive decline in Parkinson’s disease and other neurodegenerative disorders (**Fig. 3j**).^55–58^

Next, we used RNA-seq from iNs derived from healthy donors and Parkinson’s disease patients to assess age-related aspects in the resulting neurons. First, a gene set enrichment analysis was performed to determine if any molecular features relating to cellular aging were associated with donor age in iNs. Genes were ranked based on their association (using Pearson correlation coefficient) with age at sampling and six gene sets related to aging were extracted from the gene ontology database. Despite the limited age span of the donors (58 to 80 years old), we observed a positive correlation between donor age with expression of an age-related gene signature (normalized enrichment score 1.4, adjusted p-value = 0.015) (**Fig 4a-c**). To complement the GSEA data, we also looked at DNA damage, another independent marker of cellular aging, using gH2AX. This analysis comparing the number of gH2AX spots in the nucleus of parental fibroblasts with reprogrammed iNs showed a maintenance of the number of gH2AX spots after 27 days of conversion (**Fig 4d,e**). Next, we looked at the presence of the isoforms of tau that are expressed in adult mature neurons (4 repeat; 4R). This analysis showed that exon 10 (giving rise to 4R isoforms) is only expressed in iNs from adult fibroblasts (in approximately 40% of the transcripts), and not in iNs derived from fetal fibroblasts^24^ (**Fig 4f**). Moreover, the 3R/4R ratio for adult fibroblasts was 23%, whereas it was <1% for hFL1s. Taken together, this analysis suggests that donor age is at least partially maintained during iN conversion.

**Fig. 4.**
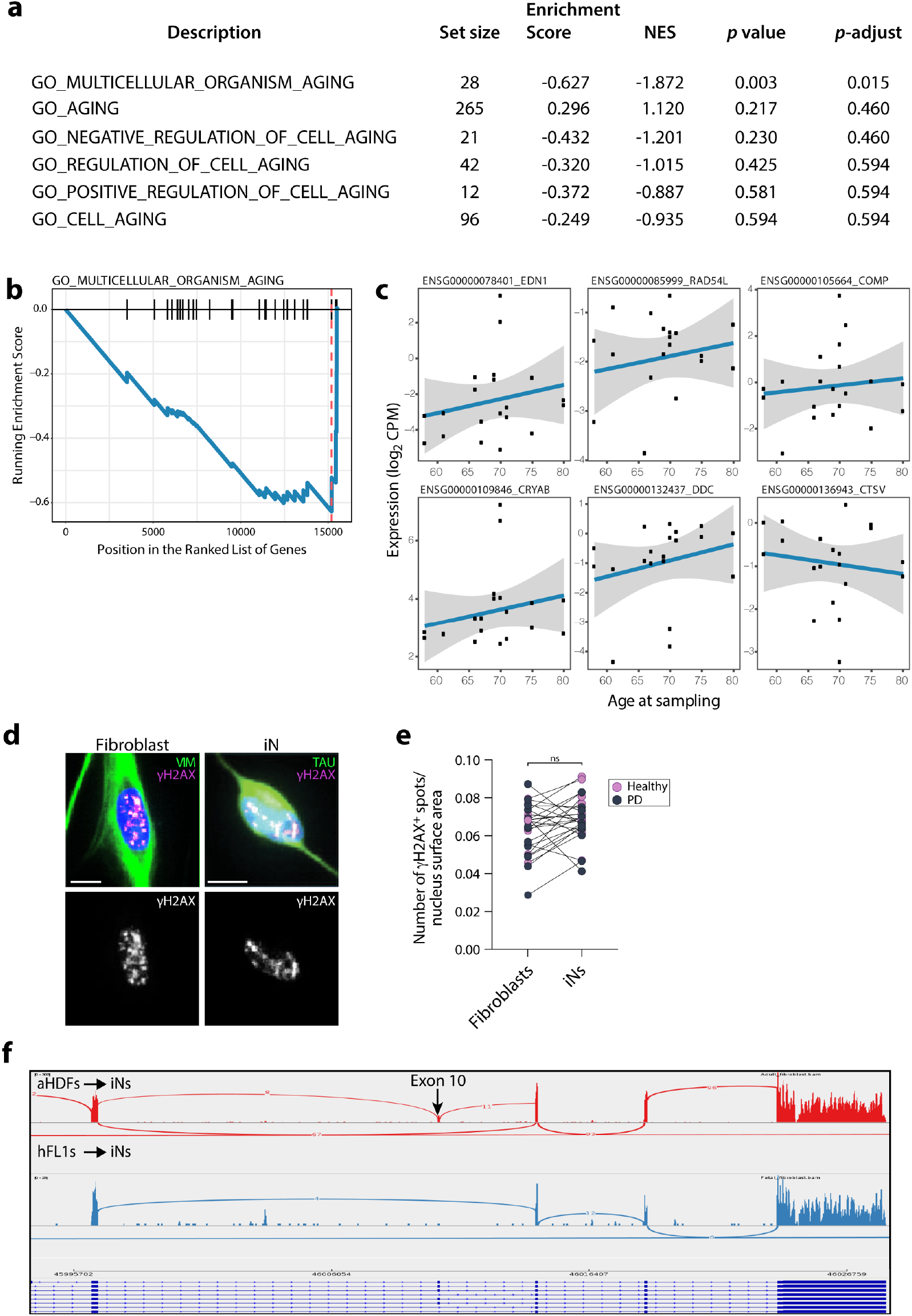
Assessment of cellular aging. **a**, Gene set enrichment analysis showing enrichment scores of pathways related to cell aging. **b**, “Multicellular organism aging” showing a significant enrichment score. **c**, After fitting a linear model adjusting for sex and disease status as covariates, genes were ranked based on their association with age at sampling. Top genes showing a clear increase in expression with age were extracted from the gene ontology database and queried against using GSEA (as implemented in the clusterProfiler R package). 5 out of 6 of these gene sets showed negative enrichment scores, indicating association of aging with donor age in this dataset. **d,** Representative image of γH2AX expression in a VIM-positive fibroblast and a TAU-positive iN, both from line #18 (87 years old). Scale bar = 10μm. e, Quantification of γH2AX-positive puncta in TAU-positive iNs (mean average of 1,327 fibroblasts and 1,210 TAU-positive cells assessed per line, n=26 lines). Two-tailed paired t-test: P=0.071, df=25. **f**, Sashimi plots visualizing splice junctions and genomic coordinates from merged bam files from adult Fib-iNs (red) and fetal Fib-iNs (blue) indicating that expression of exon 10 (4R isoforms) is only present in iNs from adult fibroblasts. Height of bars indicate expression level and the number on the lines gives number of reads spanning that splice junction. Abbreviation: ns: not significant, Vim: vimentin.

Phosphorylated αsyn is a hallmark of Parkinson’s disease pathology and this has been recapitulated in some iPSC-based cellular models of genetic forms of Parkinson’s disease^59,60^ but not idiopathic Parkinson’s disease. We therefore sought to investigate whether alterations in stress-induced autophagy in iNs from idiopathic Parkinson’s disease patients could lead to changes in the levels of phosphorylated αsyn at the Serine 129 site (pSer129 αsyn). No pSer129 αsyn staining could be detected in parental fibroblasts. However, we found that while a concurrent activation of macroautophagy by starvation and a blockage of the flux with Bafilomycin A1 did not induce significant changes in pSer129 αsyn in H-iNs (83.1 % ± 53.9 of the starved condition), it did lead to an increase in the number of PD-iNs with pSer129 αsyn-positive cytoplasmic dots (126.5 % ± 54.0 of the starved condition) (**Fig. 5a,b**). This increase in pSer129 αsyn-positive cytoplasmic dots was also observed when looking specifically in PD-iDANs as identified with TH staining (128.1% ± 21.4 of the starved condition), as compared to H-iDANs which again did not show any changes in pSer129 αsyn upon Bafilomycin A1 treatment (98.2% ± 16.0 of the starved condition) (**Supplementary Fig. 9a,b**). Finally, to assess whether elevated basal level of total αsyn could explain the elevated levels of pSer129 αsyn observed in lines starved and treated with Bafilomycin A1, we plotted the measure of the total αsyn fluorescence intensity (**Supplementary Fig. 4c**) against the pSer129 αsyn expression measured in iDANs. There was no correlation between basal total αsyn levels and the Bafilomycin A1-induced accumulation of pSer129 αsyn in iDANs (**Supplementary Fig. 9c**), suggesting that the increase seen in the Parkinson’s disease group is not due to higher basal αsyn expression.

**Fig 5.**
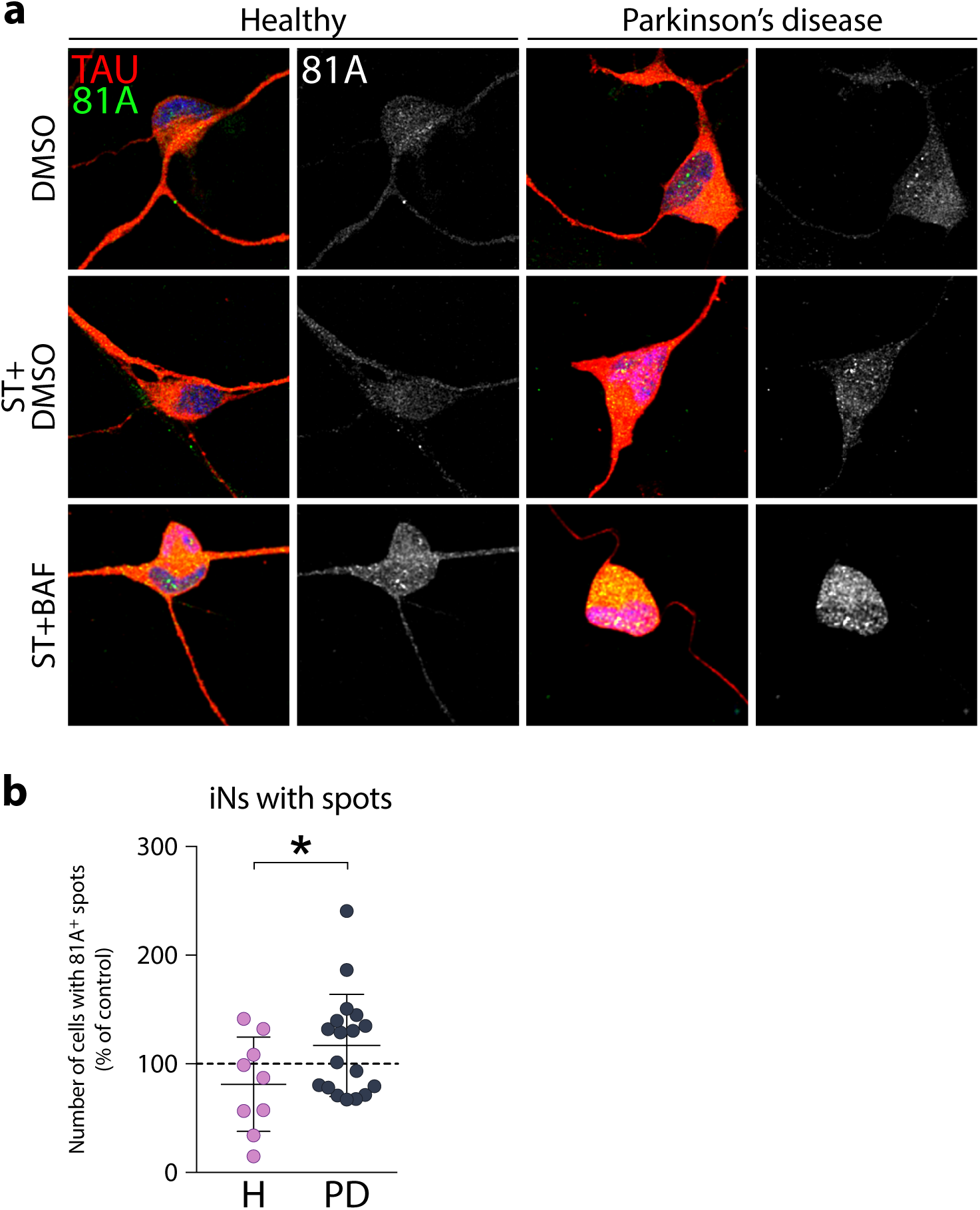
Autophagy impairments lead to an accumulation of phosphorylated asyn in PD-iNs. **a**, Confocal images of 81A-positive dot expression (αsyn pSer129) in TAU-positive iNs directly reprogrammed from fibroblasts. **b**, Quantification of TAU-positive iNs with 81A-positive puncta in the cell body (mean average of 1,461 TAU-positive cells assessed per line, n=9 healthy and n=18 Parkinson’s disease lines). Two-tailed unpaired t-test: *P=0.0329, df=25. Data were normalized as % of the control condition (% of starved+BAF/starved). Abbreviations: BAF: bafilomycin A1, H: healthy; PD: Parkinson’s disease, ST: starved.

## Discussion

This study reports age- and disease-relevant features in subtype-specific iNs derived from idiopathic Parkinson’s disease patients. Here, we have used a REST knockdown approach to enable neuronal gene transcription in adult cells,^24^ and combined that with an optimal combination of DA fate determinants (Lmx1a, Lmx1b, FoxA2, Otx2, Nurr1). This new reprogramming approach increased the efficiency, subtype identity and functional maturation of iDANs, making it possible to perform studies at a scale suitable for disease modeling, drug screening and other biomedical applications. By analyzing lines from 18 patients and 10 healthy donors, we found abnormal accumulation of structures involved in autophagy in iNs derived from idiopathic Parkinson’s disease patients, and impairment in CMA specifically in iNs with a dopaminergic phenotype. This analysis revealed that the accumulation of certain autophagic structures was restricted to neurites (e.g. accumulation of late endosomes/lysosomes). While it remains unclear as to why this occurs, such neurite specific accumulation has been previously reported in models of neurodegenerative disorders and has been attributed to deficits in retrograde transport of lysosome precursors, leading to their accumulation and further blockade in their maturation.^61^

CMA is the primary degradation pathway of the wild-type form of αsyn in neurons^46,62^ and it is present in DA neurons of the SNc.^48^ In line with this, LAMP2a and HSC70 are both down-regulated in the SNc of Parkinson’s disease patients.^63^ Our results also show a down-regulation of LAMP2a in iDANs of Parkinson’s disease patients at baseline. In addition to CMA, macroautophagy (which includes mitophagy, an autophagy pathway specific to mitochondria) has also been associated with Parkinson’s disease,^64,65^ and there is considerable crosstalk between CMA and macroautophagy. In fact, aggregates of αsyn have been shown to impair autophagosome clearance during macroautophagy.^66^ Our study shows that by blocking the autophagic flux the context of in stress-induced autophagy, an accumulation of pSer129 αsyn in PD-iNs/iDANs was seen, but not in iNs/iDANs from control donors.

The phosphorylation of αsyn at Serine 129 is observed in the brain of Parkinson’s disease patients^67^ and is thought to be critical in Parkinson’s disease pathogenesis.^68^ Moreover, aggregate clearance of αsyn depends mainly on the autophagy pathway,^69^ which is supported by the data in our study showing an increase in pSer129 αsyn in PD-iNs/iDANs. However, one outstanding question is whether the accumulation of pSer129 αsyn is triggered by, or modulated by, similar factors in every case of idiopathic Parkinson’s disease. Arguments against this hypothesis include that i) at the cellular level, other pathophysiological processes have been identified in models and post-mortem Parkinson’s disease brains that lead to pSer129 αsyn accumulation, and ii) it is increasingly recognized that idiopathic Parkinson’s disease can be stratified based on different parameters, including clinical features and rate of progression and with this, extent of pathology.^70^ As such, cellular processes that trigger pSer129 αsyn accumulation are likely to differ within the idiopathic Parkinson’s disease patient population, leading to the clinical heterogeneity observed. In order to study such a hypothesis, there is a great need for models of idiopathic Parkinson’s disease that can reveal such differences in Parkinson’s disease-associated pathophysiology.

In the study we report here, a new reprogramming method allowed us to compare iNs from 18 different idiopathic Parkinson’s disease patients that were all processed in parallel, and which uniquely revealed baseline disease-relevant pathology in the neurons to some extent across all the lines. We observed alterations in stress-induced autophagy as a group effect as compared to sex- and age-matched healthy donor lines. Disease-associated impairment could not be detected in the parental fibroblasts. This shows that direct conversion of fibroblasts, where age-related aspects of the donor is maintained, uniquely allows for cell-based models of idiopathic Parkinson’s disease for which iPSC-based modeling is challenging due to the large number of patient lines that needs to be studied and the inability to generate isogenic controls. Importantly, our cellular model based on fibroblast-to-neuron conversion showed that iNs from different patients are not impaired to the same degree, and as such, the iDAN system is more likely suitable to molecular-based stratification of idiopathic Parkinson’s disease while iPSC-derived neurons from patients with monogenic Parkinson’s disease and their isogenic controls may be more suitable to study more defined aspects of disease pathology. In line with the use of iDANs for disease stratification, we found that the degree of impairment relates, at least to some extent, to the age of the donor, the age of onset of their Parkinson’s disease and their Tau haplotype. This effect of age and genetic variance on disease pathology has not been modeled before and suggests that direct conversion to iDANs could be used for differential diagnostics, drug screening and disease modeling of late onset neurogenerative diseases.

On a more general level, our results demonstrate the importance of establishing models of neurodegenerative disease with cells that resemble the subtype and functionality of the affected neurons in individual patients as closely as possible. For example, we could not detect any autophagy-related impairment in the fibroblasts prior to conversion, clearly demonstrating that the reprogramming to neurons is essential to reveal disease-related phenotypes, especially as some phenotypes were detected in neurites. Also, specific CMA impairments were detected only in iNs with a DA phenotype. In addition to this, our data supports other studies reporting that direct reprogramming maintains the aging signature of the donor cell,^6,15–18^ which uniquely allows for modeling aspects of disease such as age of onset.

Taken together, our study demonstrates the potential of direct neuronal conversion to study late onset neurodegenerative diseases. It is the first study to model disease heterogeneity within the idiopathic Parkinson’s disease population at the neuronal level in directly converted cells, and to report that stress-induced macroautophagy impairments are present and linked to the accumulation of relevant αsyn pathology in the cells. Future studies using this cellular model will contribute to a deeper understanding of the age-associated pathology of Parkinson’s disease along with the cellular basis of disease subtypes and variable progression and through this allow us to better assess therapeutic interventions.

## Materials and methods

### Cell lines, genotyping and DNA-sequencing

Adult dermal fibroblasts were obtained from the Parkinson’s Disease Research clinic at the John van Geest Centre for Brain Repair (Cambridge, UK) and used under full local ethical approvals: REC 09/H0311/88 (University of Cambridge) and CERSES-18-004-D (University of Montreal) (**Table 1**). The subjects’ consent was obtained according to the declaration of Helsinki. Cell lines used in this study will be available subject to appropriate ethical approval and an MTA from the requestor. For biopsy sampling information see.^24^ All patients were screened for three common Mendelian mutations associated with late onset Parkinson’s Disease: LRRK2 G2019S as well as GBA L444P and GBA N370S. One out of 19 patients was identified as a LRRK2 G2019S mutation carrier and was removed from further analysis. Samples were also screened for SNCA gene expression levels in our RNA-seq dataset to detect any overexpression of the SNCA gene in the iN samples that would suggest a duplication or triplication. All Parkinson’s Disease samples included in the RNA-seq analysis had SNCA expression that followed a normal distribution without any outliers. For MAPT haplotype genotyping, single nucleotide polymorphism (SNP) genotyping was undertaken using a predesigned assay, rs9468 (Applied Biosystems), tagging the MAPT H1 versus H2 haplotype, and run on a Quantistudio 7 Flex Real-Time PCR System (ThermoFisher), according to the manufacturer’s instructions. There were no inconsistencies amongst the 28 samples genotyped in triplicates.

### Cell culture

Fibroblasts were expanded in T75 flasks with standard fibroblast medium (DMEM, 10 % FBS, 100U/mL penicillin-streptomycin) at 37°C in 5 % CO_2_. After thawing, cells were kept for a minimum of two days in culture before starting experiments. When confluent, the cells were dissociated with 0.05 % trypsin and plated at a lower density to expand them. To freeze the fibroblasts from a confluent T75 flask, the cells were detached after 5 minutes incubation in 0.05 % trypsin at 37°C, spun for 5 minutes at 400 g and frozen in a 50/50 mixture of DMEM and FBS with 10 % DMSO. All cell lines were routinely tested for mycoplasma and were negative. For experiments on fibroblasts, cells were plated at a density of 1,900-3,800 cells per cm^2^ in 24-well plates (Nunc) and analyses were performed three days later.

### Viral vectors and virus transduction

DNA plasmids expressing mouse open reading frames (ORFs) for *Ascl1, Lmx1a, Lmx1b, FoxA2, Otx2, Nurr1, Smarca1, CNPY, En1 or Pax8* in a third-generation lentiviral vector containing a non-regulated ubiquitous phosphoglycerate kinase (PGK) promoter were generated, as well as two short hairpin RNAs (shRNAs) targeting RE1-silencing Transcription Factor (REST) containing a non-regulated U6 promoter. Plasmids used in this study have been deposited in Addgene (#33013, #33014, #34997, #35000, #35001, #127573, #127574) Furthermore, the pB.pA.shREST all-in-one vector from^24–26^ was used to reprogram iNs for RNAseq and western blot (WB). All the constructs have been verified by sequencing. Lentiviral vectors were produced as previously described^27^ and titrated by qPCR analysis.^28^ Transduction was performed at a MOI of 5 for each vector (all viruses used in this study were titered between 1 x 10^8^ and 9 x 10^9^) or MOI of 20 in the case of the pB.pA.shREST vector.

### Neural reprogramming

For direct neural reprogramming, fibroblasts were plated at a density of 26,300 cells per cm^2^ in 24-well plates (Nunc). Prior to plating, the wells were coated overnight with either 0.1 % gelatin (Sigma), or a combination of PFL for long-term cultures: Polyornithine (15 μg/mL), Fibronectin (0.5 ng/μL) and Laminin (5 μg/mL). Cells used for electrophysiological recordings were directly plated onto glass coverslips coated with PFL as described in^26^. Three days after the viral transduction, the fibroblast medium was replaced with early neural differentiation medium (ENM) (NDiff227; Takara-Clontech) supplemented with growth factors at the following concentrations: LM-22A4 (2 μM, R&D Systems), GDNF (2 ng/mL, R&D Systems), NT3 (10 ng/μL, R&D Systems), as well as with db-cAMP (0.5 mM, Sigma) and the small molecules CHIR99021 (2 μM, Axon), SB-431542 (10 μM, Axon), noggin (0.5 μg/ml, R&D Systems), LDN-193189 (0.5 μM, Axon), valproic acid sodium salt (VPA; 1mM, Merck Millipore). Half medium changes were performed twice a week. At 18 days post-transduction, the small molecules were stopped, and the neuronal medium was supplemented only with LM-22A4 (2 μM), GDNF (2 ng/mL), NT3 (10 ng/μL) and db-cAMP (0.5 mM) until the end of the experiment (Late neuronal medium; LNM). To assess the reprogramming efficiency of each line, all 28 lines were reprogrammed at the same time with the same virus mixture, and this was repeated three times using different batches of virus for each of the 8 lentiviral vectors required for the iDAN reprogramming.

### Whole cell patch clamp recordings

Prior to recording, the cells on the coverslips were transferred from the culture medium to BrainPhys medium ^29^ for 30 minutes and maintained at 34.5 °C. Cells were then moved to a recording chamber and submerged in a flowing artificial cerebrospinal fluid (ACSF) solution gassed with 95 % O_2_ and 5 % CO_2_. The composition of the ACSF was (in mM): 126 NaCl, 2.5 KCl, 1.2 NaH_2_PO_4_-H_2_O, 1.3 MgCl_2_-6H_2_O, and 2.4 CaCl_2_-6H_2_O, 22 NaHCO_3_, 10 glucose adjusted to pH = 7.4. Temperature of the chamber was maintained at 34 °C throughout the entire recording session. Multi-clamp 700B (Molecular Devices) was used for the recordings and signals were acquired at 10 kHz using pClamp10 software and a data acquisition unit (Digidata 1440A, Molecular Devices). Current was filtered at 0.1 Hz and digitized at 2 kHz. Borosilicate glass pipettes ranging between 4-7 MΩ were used and they were filled with the following intracellular solution (in mM): 122.5 potassium gluconate, 12.5 KCl, 0.2 EGTA, 10 Hepes, 2 MgATP, 0.3 Na_3_GTP and 8 NaCl adjusted to pH = 7.3 with KOH as in^11^. The intracellular solution was kept on ice during the recordings. Cells with neuronal morphology characterized by a rounded cell body were selected for recordings. Input resistances and injected currents were monitored throughout the experiments. Passive properties of the membrane were monitored, and recordings were discarded when changes in the capacitance were higher than 20 % from the beginning to the end of the recording session. Resting membrane potentials were monitored immediately after breaking-in, in current-clamp mode. The membrane potential was kept between −40 mV to −60 mV and currents were injected for 500 ms from −20 pA to +90 pA with 10 pA increments to induce action potentials. The number of action potentials for each cell was taken as the highest frequency of action potential induced by a step of current within the same cell and averaged over the total cells patched per line. Voltage ramp was characterized by constant increase in voltage from −70 mV to +20 mV in 0.5 sec intervals. Inward sodium and delayed rectifying potassium currents were measured in voltage clamp at depolarizing steps of 10 mV for 100 ms. Spontaneous firing was recorded in voltage-clamp mode at resting membrane potentials.

### Starvation and Bafilomycin A1 treatment

On day 28 following viral transduction, iNs were starved for 4 hours by replacing the culture medium with HBSS and Ca^2+^/Mg^2+^ and compared to the condition without starvation, where cells were left in their original culture medium. The duration of starvation treatment was chosen based on a starvation curve performed on the iNs (0, 2, 4h), which showed clear increases in p62 and microtubule-associated protein 1 light chain 3 beta, LC3) expression by WB (**Supplementary Fig. 2b,c**) in the absence of neuronal cell death (**Supplementary Fig. 2a**). For the experiment with Bafilomycin A1, cells were starved in HBSS Ca^2+^/Mg^2+^ containing Bafilomycin A1 (100 nM; Sigma Aldrich) for 2 hours and compared to cells incubated in HBSS Ca^2+^/Mg^2+^ containing dimethyl sulfoxide (DMSO; vehicle). This regimen was chosen based on the increase of LC3-II and the LC3-II/LC3-I ratio as assessed by WB (**Supplementary Fig. 2d,e**). At the end of the incubation period, cells were fixed in 4 % paraformaldehyde.

### Western blots

Cells were lysed and homogenized as described elsewhere.^30^ Protein concentration was determined using a DC protein assay kit (Bio-Rad, 5000116). 10-15 μg of protein was boiled at 95°C for 5 min in Laemmli buffer (Bio-Rad), separated on a 4–12 % SDS/PAGE gel and then transferred using the Transblot®-Turbo™ Transfer system (Bio-Rad). After 1 hour blocking in Tris-buffered saline (TBS; 50 mM Tris-Cl, 150 mM NaCl, pH 7.6) with 0.1 % Tween 20 (Sigma-Aldrich, P7949) and 2.5 % (wt:vol) non-fat dry milk (Bio-Rad Laboratories), membranes were incubated overnight at 4°C in one of the primary antibodies summarized in **Supplementary Table 1**. After washing with TBST, membranes were incubated for 1 hour at room temperature with HRP-conjugated secondary antibodies. Protein expression was developed with the ECL™ Prime Western Blotting Detection Reagent (Life Technologies, RPN2232). Signal was captured using a Chemidoc MP system (Bio-Rad). Band intensity was quantified using ImageJ software (ImageJ, 1.48v) by densitometry.

### Immunocytochemistry and high content screening quantifications

Following fixation in 4 % paraformaldehyde, cells were permeabilized with 0.1 % Triton-X-100 in 0.1 M PBS for 10 minutes. Thereafter, cells were blocked for 30 minutes in a solution containing 5 % normal serum in 0.1 M PBS. The primary antibodies used are listed in **Supplementary Table 1** and were diluted in the blocking solution and applied overnight at 4 °C. Fluorophore-conjugated secondary antibodies (1:200; Jackson ImmunoResearch Laboratories) as well as 4’,6-diamidino-2-phenylindole (DAPI; 1:1,000, Sigma Aldrich) were diluted in blocking solution and applied for 2 hours. Fluorescence images of the Heat shock cognate 71 kDa protein (HSC70, also known as HSPA8), lysosome-associated membrane protein 2a (LAMP2a) and tyrosine hydroxylase (TH) and the TAU-81A stainings were taken using a confocal laser scanning microscope (Leica, TCS SP8), whereas the rest of the images were taken using either an inverted microscope (Leica, DFC360 FX-DMI 6000B) or CellInsight CX5 or CX7 High-Content Screening (HCS) microscopes (Thermo Scientific).

The total number of DAPI-positive, TAU-positive and TH-positive cells per well, as well as the average fluorescence intensity for αsyn was quantified using the Cellomics Array Scan (Array Scan VTI, Thermo Fischer), which is an automated process ensuring unbiased measurements between groups. Applying the program “Target Activation”, 100-200 fields (10 X magnification) were acquired in a spiral fashion starting from the center. The same array, run at 20 X magnification, was used for the analysis of the number of neurites per TAU-positive cell using the program “Neuronal Profiling”. Neuronal purity was calculated as the number of TAU- or MAP2-positive cells over the total number of cells in the well at the end of the experiment. Dopaminergic subtype purity was calculated as the number of TH- or aldehyde dehydrogenase 1 family member A1 (ALDH1A1)-positive cells over the total number of TAU- or MAP2-positive cells in the well at the end of the experiment. Average dot number and size was measured in those neurons in which the cytoplasm and neurites were defined by TAU or TH staining. Puncta of p62, LC3, LAMP2, LAMP2a, HSC70, αsyn and 81A were detected (using a “Spot Detection” program) and measured in each case. Experiments done on fibroblasts were quantified using the same approach, with quantification of puncta of p62, LC3, LAMP2, LAMP2a and HSC70 in the cytoplasm, defined by vimentin (VIM)-positive staining. For gH2AX measurements, puncta positive for gH2AX were detected in the nuclei of fibroblasts and iDANs (defined by the DAPI stained region) at 20 X magnification using a “Spot Detection” program.

### qRT-PCR for neuronal gene expression

Total RNA was extracted from human fibroblasts as well as iNs from the same lines using the miRNeasy kit (Qiagen) followed by Universal cDNA synthesis kit (Fermentas). Three reference genes were used for each qPCR analysis (ACTB, GAPDH and HPRT1). All primers were used together with LightCycler 480 SYBR Green I Master (Roche). Standard procedures of qRT-PCR were used, and data quantified using the ΔΔCt-method. Statistical analyses were performed on triplicates. A custom RT^2^ profiler PCR Array (Qiagen) containing 90 neuronal genes was also used according to the manufacturer instructions.

### RNA-seq analysis

Fibroblasts from the healthy donor (n = 10) and Parkinson’s disease (n = 10) lines were plated and either collected for RNA extraction following 3 days in culture or transduced with the pB.pA.shREST lentiviral vector^24–26^ the day following plating and allowed to be reprogrammed for 30 days. RNA was extracted using the RNeasy mini kit (Qiagen) with DNase treatment. cDNA libraries were prepared using the Illumina truseq library preparation kit and sequenced with 2 x 150 bp paired end reads on an Illumina NextSeq 500 High Output kit. Raw base calls were demultiplexed and converted into sample specific fastq format files using default parameters of the bcl2fastq program provided by illumina. Quality of reads was checked using FastQC and multiQC tools. Reads were mapped to the human genome (GRCh37) using the STAR mapping algorithm.^31^ mRNA expression was quantified using RSEM^32^ and Ensembl version 75 as the gene model. Read counts were normalized to the total number of reads mapping to the genome. After sample level QC, differential expression analysis was done using the limma/voom algorithm.^33^ Downstream analyses were performed using in house R scripts. Gene Set Enrichment Analysis was performed by fitting a linear model adjusting for sex and disease status as covariates - genes were then ranked based on their association with age at sampling. Six gene sets related to aging were extracted from the gene ontology database and queried using gene set enrichment analysis (GSEA) (as implemented in the clusterProfiler R package). Differential splicing of the MAPT gene was visualized in IGV (version 2.8). Significant up and down regulated pathways were selected using Bonferroni post hoc corrected p values (padj < 1e-4).

### Statistical analysis

All data are expressed as mean ± the standard deviation. Whenever the analysis is performed with one cell line, biological replicates (*n* = 3-4) were used. In case of experiments using multiple cell lines, we used a minimum of *n* = 5 to account for inter-individual variation. A Shapiro-Wilk normality test was used to assess the normality of the distribution. When a normal distribution could not be assumed, a non-parametric test was performed. Groups were compared using a one-way ANOVA with a Bonferroni post hoc or a Kruskal-Wallis test with a Dunn’s multiple comparisons tests. To determine whether there was a significant difference between two sets of observations repeated on the same lines, a paired sample t-test was also performed. In case of only two groups, they were compared using a Student *t*-test. An F test was used to compare variance and in case of unequal variance a Welch’s correction test was then performed. Statistical analyses were conducted using the GraphPad Prism 8.0. An alpha level of p < 0.05 was set for significance.

## Data availability

The RNAseq dataset can be found on the GEO repository under accession number GSE125239.

## Acknowledgements

We thank Marie Persson Vejgården, Sol Da Rocha Baez, Ulla Jarl (Lund University), and Dr. Maria Ban (Neurology Unit at the University of Cambridge) for technical assistance as well as Dr. Anna Hammarberg at the MultiPark Cellomics platform at Lund University for her valuable help with high content screening.

## Abbreviations

αsyn: alpha-synuclein
ALDH1A1: aldehyde dehydrogenase 1 family member A1
ACSF: artificial cerebrospinal fluid
CMA: chaperone-mediated autophagy
DAPI: 4’,6-diamidino-2-phenylindole
DMSO: dimethyl sulfoxide
GSEA: gene set enrichment analysis
ENM: early neural differentiation medium
DA: dopaminergic
HSC70: Heat shock cognate 71 kDa protein
iDANs: induced DA neurons
iNs: induced neurons
iPSCs: induced pluripotent stem cells
LAMP2: lysosome-associated membrane protein 2
LC3: microtubule-associated protein 1 light chain 3 beta
LNM: Late neuronal medium
PGK: phosphoglycerate kinase
REST: RE1-silencing Transcription Factor
TTX: tetrodotoxin
TH: tyrosine hydroxylase
VIM: vimentin
WB: western blot

## Supplementary Material

**Supplementary Fig. 1.**
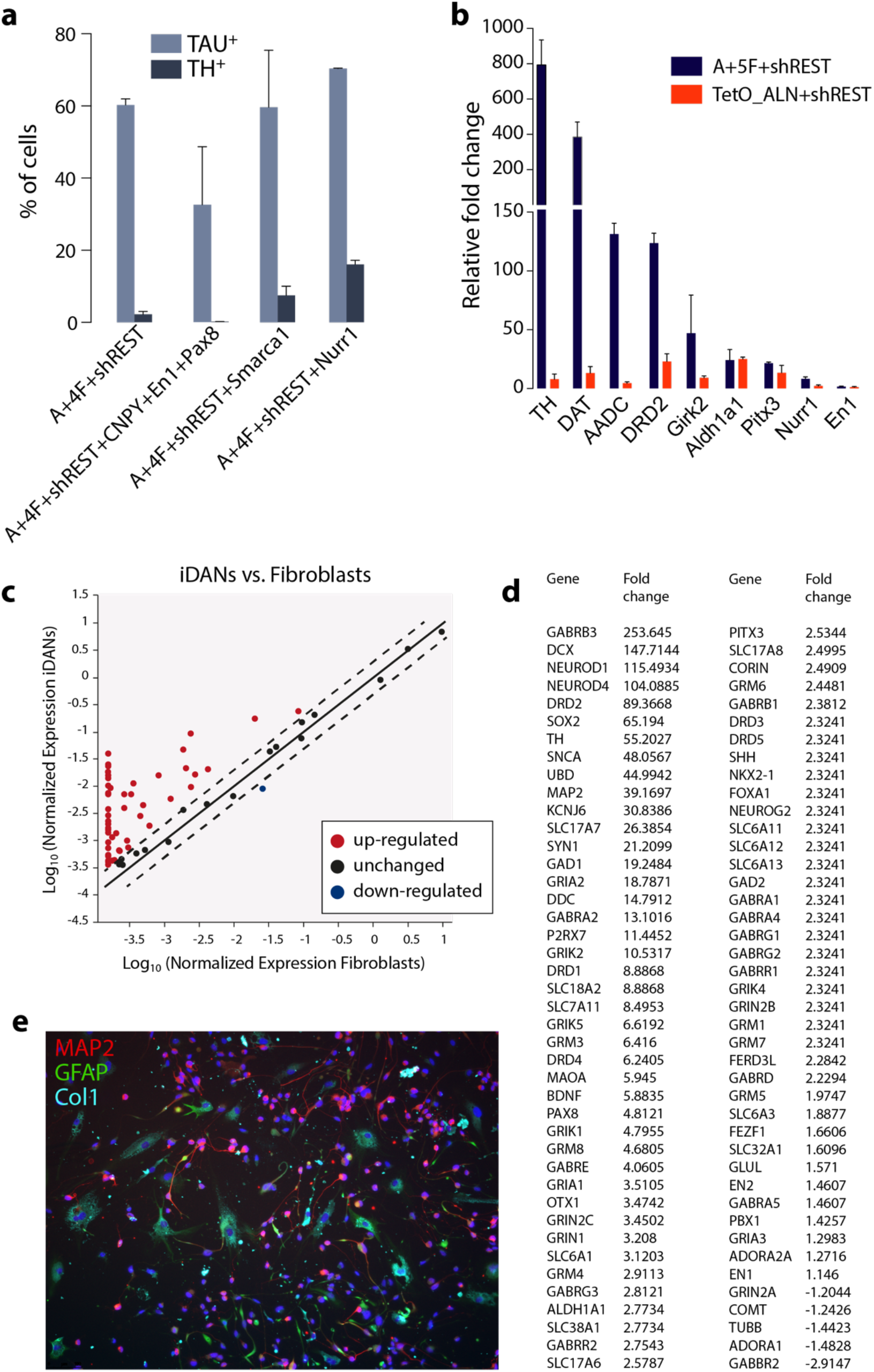
Generation of iDANs. **a**, Quantification of TAU-positive and TH-positive cells using different reprogramming factors. 4F: Lmx1a, Lmx1b, FoxA2, Otx2. **b,** Gene expression quantification of DA genes relative to parental fibroblast levels (from 2 to 3 biological replicates). 5F: Lmx1a, Lmx1b, FoxA2, Otx2, Nurr1. TetO_ALN: Ascl1, Lmx1a, Nurr1 from Caiazzo et al. 2011 ^8^. **c**, Fold change of all of the neuronal and dopaminergic related genes on the qPCR array as compared to parental fibroblasts. Significantly up- and down-regulated genes are in red and blue, respectively. **d**, Fold change of the top up-regulated neuronal, dopaminergic, glutamatergic and GABAergic related genes as compared to parental fibroblasts. **e**, MAP2, GFAP and Collagen 1 expression at 25 days following the start of conversion with the pB.pA.shREST all-in-one vector.

**Supplementary Fig. 2.**
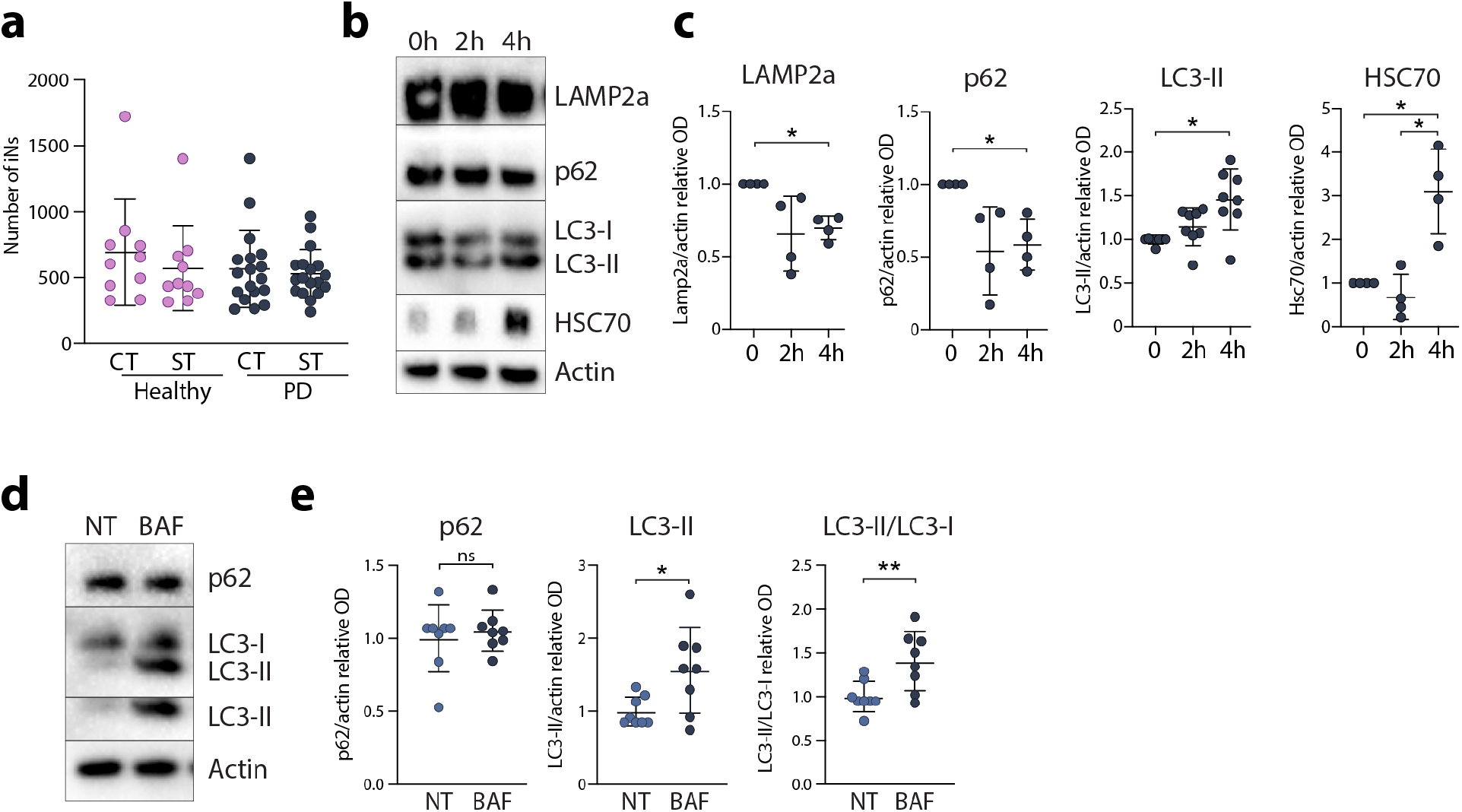
Validation of starvation and Bafilomycin regimen in iNs. **a**, Quantification of the number of TAU+ iNs following a 4-hour starvation period. **b**, Representative WB image showing LAMP2a, SQSTM1 (p62), LC3, HSC70 and Actin immunolabeling in iNs from a healthy donor starved for 0, 2 and 4 hours. **c**, OD quantification of LAMP2a, SQSTM1 (p62), LC3, HSC70 immunoblots in iNs starved for 0, 2 and 4 hours. One-way ANOVA on repeated measures, Tukey post-hoc: *p<0.05. Data are shown as mean ± SD. Values were normalized to non-treated expression levels and corrected to actin values. **d**, Representative WB image showing SQSTM1 (p62), LC3 and Actin immunolabeling in non-treated (NT) and Bafilomycin A1 (BAF) treated healthy donor iNs. **e**, OD quantification of p62, LC3 immunoblots in NT and BAF treated healthy donor iNs. Two-tailed unpaired t-test: *P<0.05; **P<0.01. Data are shown as mean ± SD. Values were normalized to non-treated expression levels and corrected to actin values.

**Supplementary Fig. 3.**
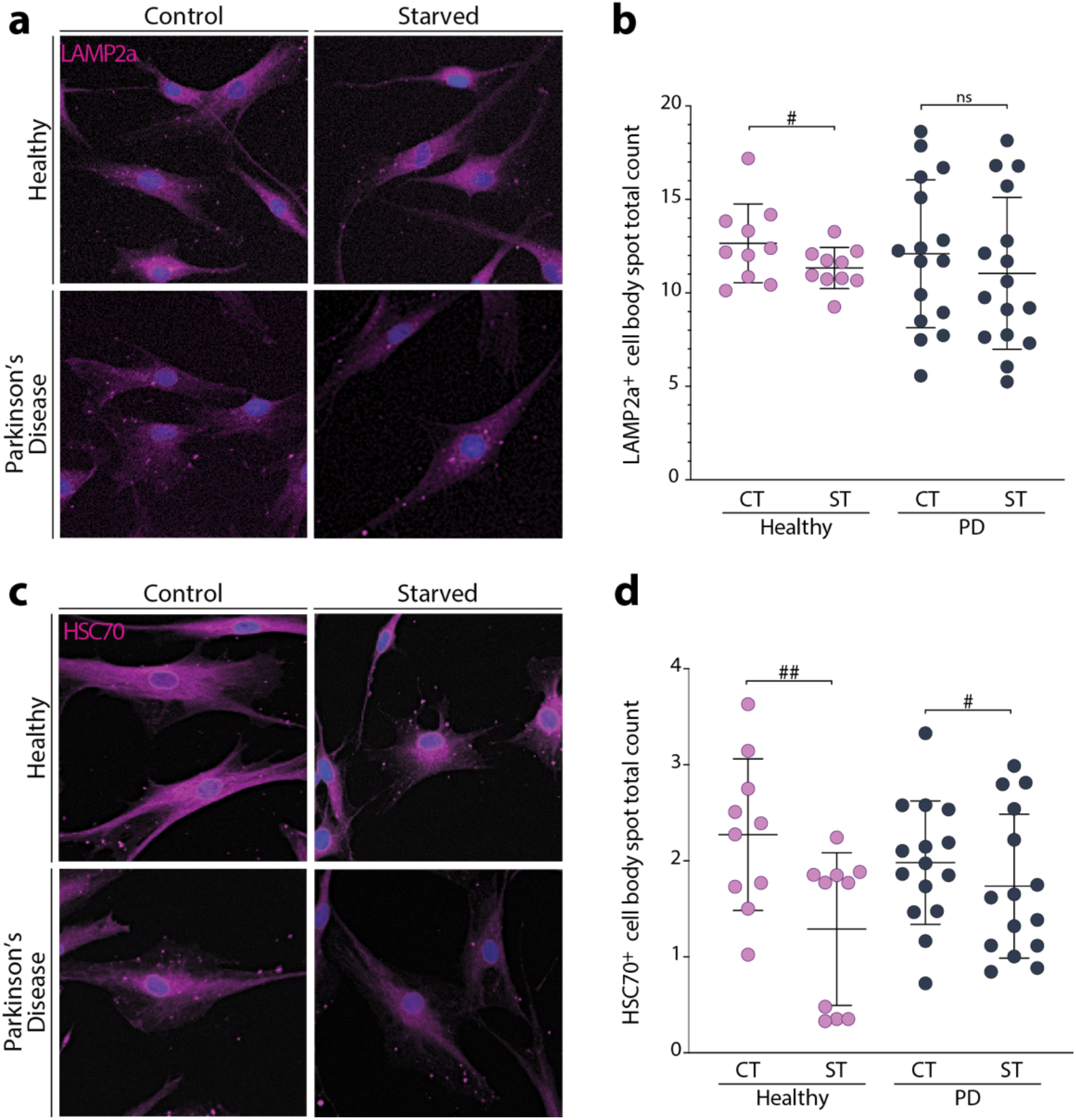
No accumulation of LAMP2a and HSC70 in PD-Fibroblasts upon starvation. **a**, LAMP2a-positive dot expression in fibroblasts. **b**, Quantification of LAMP2a-positive dots in fibroblasts (mean average of 2,949 cells assessed per line). One-way ANOVA, Bonferroni post-hoc: P=0.59. Paired student’s t-test: #P<0.05, as compared to the non-starved condition. **c**, HSC70-positive dot expression and spot detection analysis in fibroblasts. **d**, Quantification of HSC70-positive dots in fibroblasts (mean average of 2,949 cells assessed per line). Kruskal-Wallis test, Dunn’s multiple comparisons test: P=0.11; H: Wilcoxon matched pairs signed rank test: ^##^P=0.0098, rs=0.346. PD: Two-tailed paired t-test: ^#^P=0.0406, df=13. Abbreviations: CT: control, H: healthy, ns: not significant, PD: Parkinson’s disease, ST: starved.

**Supplementary Fig. 4.**
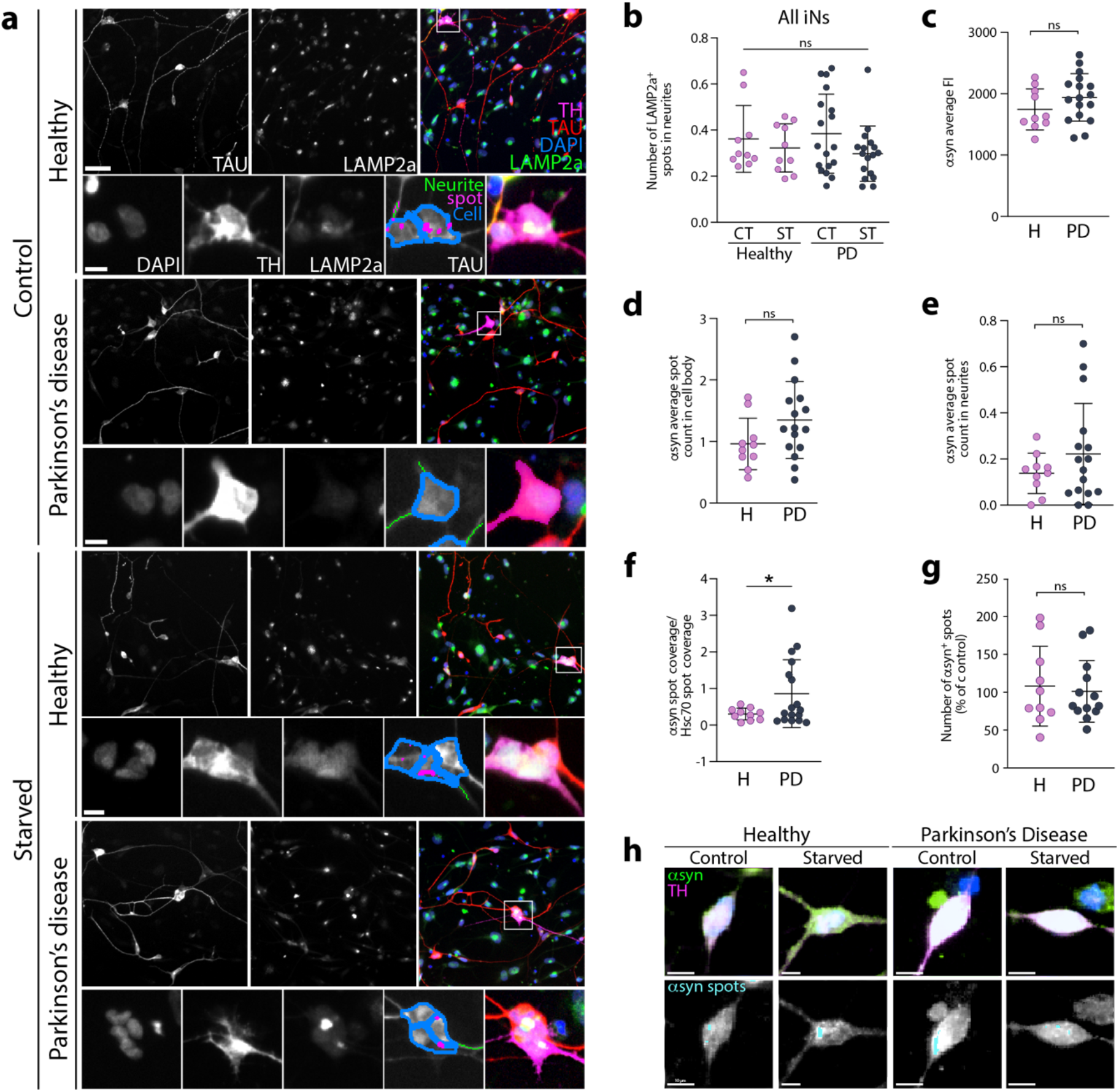
Chaperone-mediated autophagy impairment in PD-iDANs. **a**, LAMP2a-positive dot expression and spot detection analysis of LAMP2a-positive dots (in pink) in double TAU-positive/TH-positive iDANs. Scale bar = 50μm; insets = 10μm. **b**, Quantification of LAMP2a-positive dots in neurites of TAU-positive iNs (mean average of 603 iNs assessed per line). **c**, Quantification of αsyn-positive fluorescence intensity in TH-positive iDANs. **d**, Quantification of αsyn-positive spot count in the cell body of TH-positive iDANs. **e**, Quantification of αsyn-positive spot count in the neurites of TH-positive iDANs. **f**, Quantification of αsyn-positive/HSC70 spot coverage in the cell body of TH-positive iDANs. Two-tailed unpaired t-test with Welch’s correction: *P=0.0267, df=17.62. **g**, Quantification of αsyn-positive spot count in TH-positive iDANs following starvation. **h**, αsyn-positive dot expression and spot detection analysis of αsyn-positive dots (in cyan) in TH-positive iDANs. Scale bar = 10μm. Abbreviations: ns: not significant.

**Supplementary Fig. 5.**
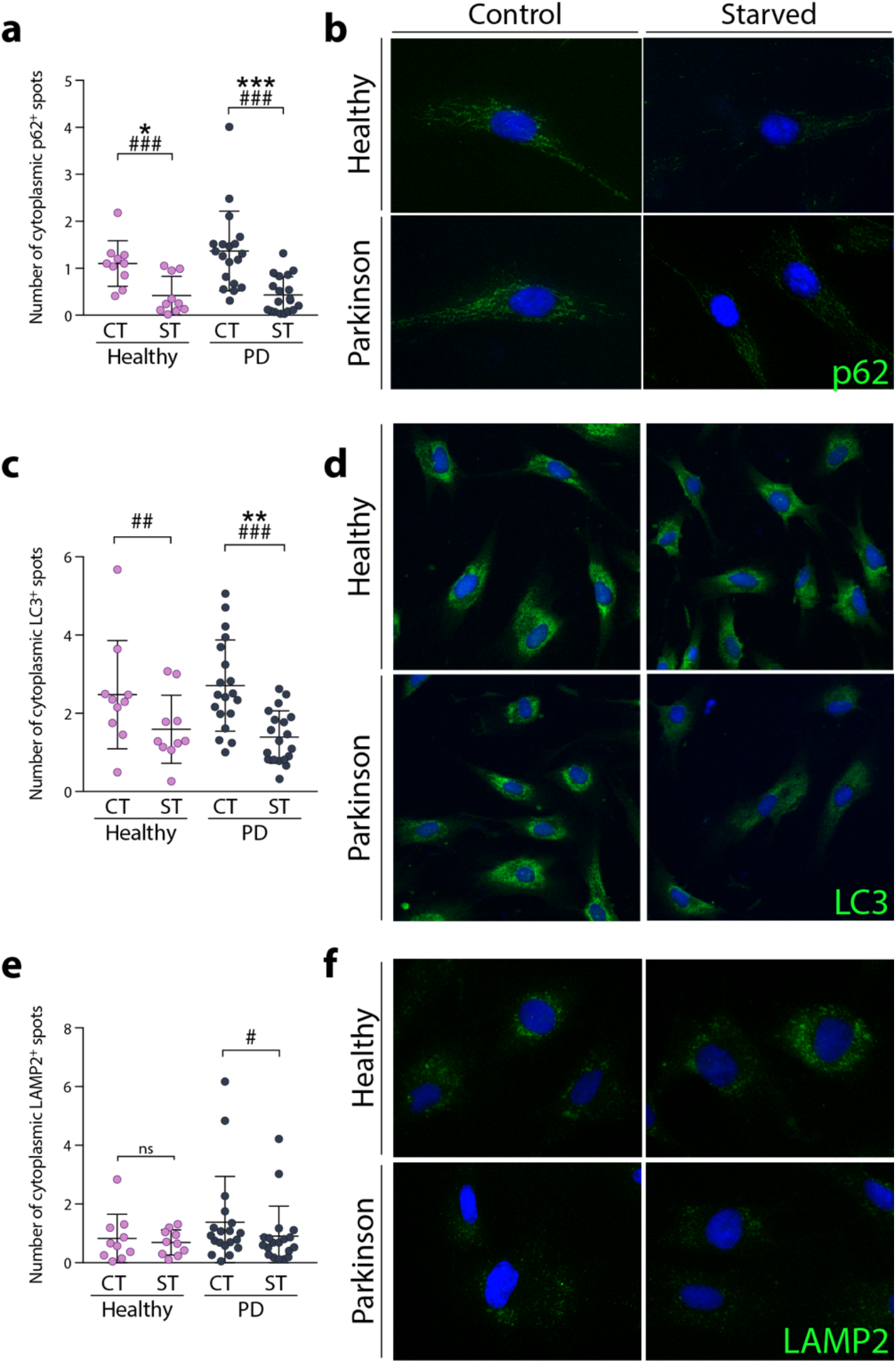
No accumulation of p62, LC3 and LAMP2 in PD-Fibroblasts upon starvation. **a**, p62-positive dot expression and spot detection analysis in fibroblasts. **b**, Quantification of p62-positive dots in fibroblasts (mean average of 625 cells assessed per line). One-way ANOVA, Bonferroni post-hoc: *p<0.05, ***p<0.001. Paired student’s t-test: ###p<0.001 as compared to the condition without starvation. **c**, LC3-positive dot expression and spot detection analysis in fibroblasts. **d**, Quantification of p62-positive dots in fibroblasts (mean average of 651 cells assessed per line). One-way ANOVA, Bonferroni post-hoc: **p<0.01, as compared to the healthy group. Paired student’s t-test: ##p<0.01, ###p<0.001 as compared to the condition without starvation. **e**, LAMP2-positive dot expression and spot detection analysis in fibroblasts. **f**, Quantification of LAMP2-positive dots in fibroblasts (mean average of 1,126 cells assessed per line). Paired student’s t-test: #p<0.05 as compared to the condition without starvation. Abbreviations: CT: control, H: healthy, ns: not significant, PD: Parkinson’s disease, ST: starved.

**Supplementary Fig. 6.**
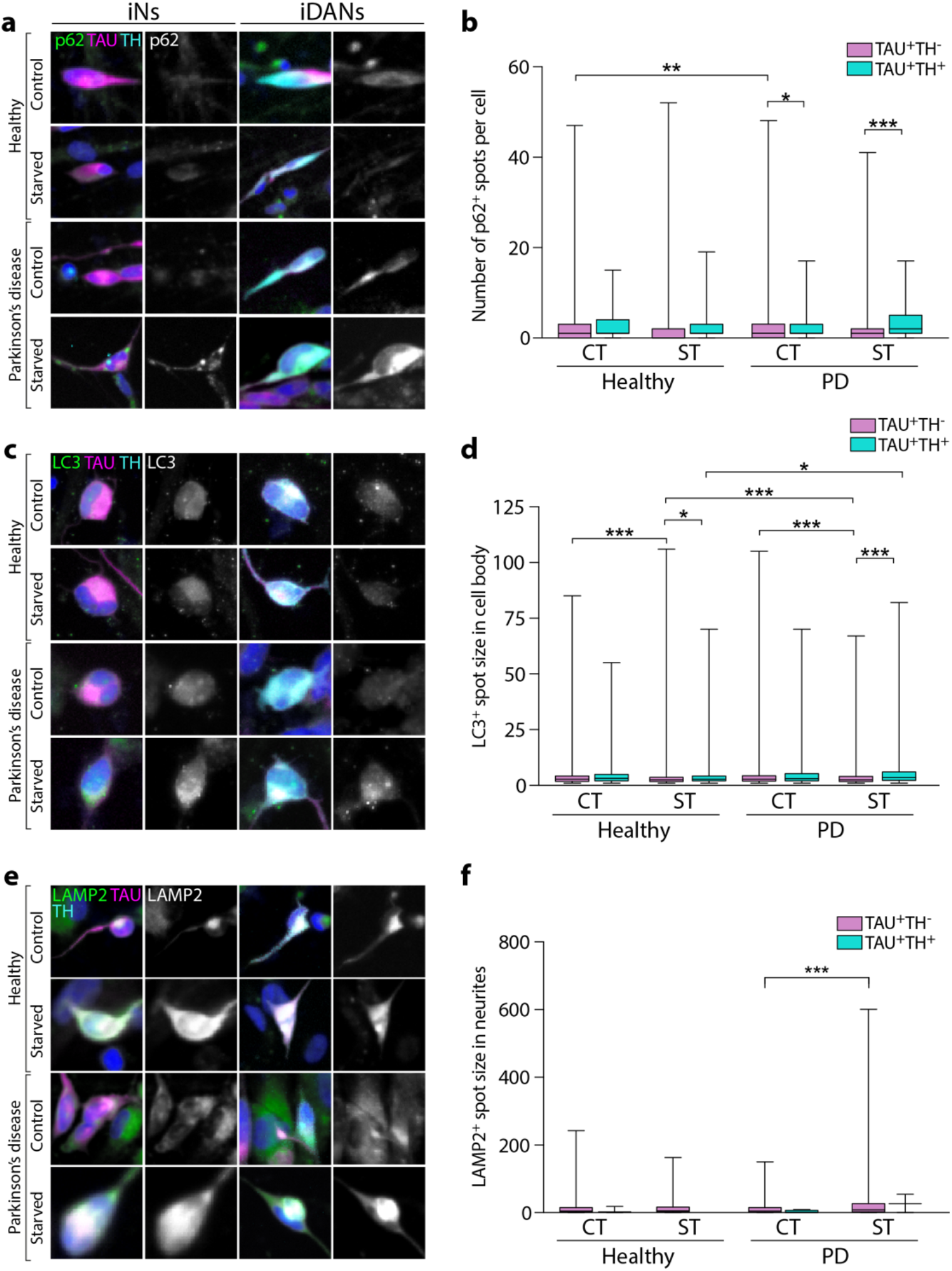
p62, LC3 and LAMP2 in iNs vs. iDANs upon starvation. **a**, p62-positive dot expression in TAU-positive/TH-negative and TAU-positive/TH-positive cells. **b**, Whisker plot showing quantification of p62-positive dots in TAU-positive /TH-negative and TAU-positive /TH-positive cells. Kruskal-Wallis test, Dunn’s multiple comparisons test: *p<0.05, **p<0.01, ***p<0.001. **c**, LC3-positive dot expression in TAU-positive/TH-negative and TAU-positive/TH-positive cells. **d**, Whisker plot showing quantification of LC3-positive dots in TAU-positive/TH-negative and TAU-positive/TH-positive cells. Kruskal-Wallis test, Dunn’s multiple comparisons test: *p<0.05, ***p<0.001. **e**, LAMP2-positive dot expression in TAU-positive/TH-negative and TAU-positive/TH-positive cells. **f**, Whisker plot showing quantification of LAMP2-positive dots in TAU-positive/TH-negative and TAU-positive/TH-positive cells. Kruskal-Wallis test, Dunn’s multiple comparisons test: ***p<0.001. Abbreviations: CT: control, H: healthy, PD: Parkinson’s disease, ST: starved.

**Supplementary Fig. 7.**
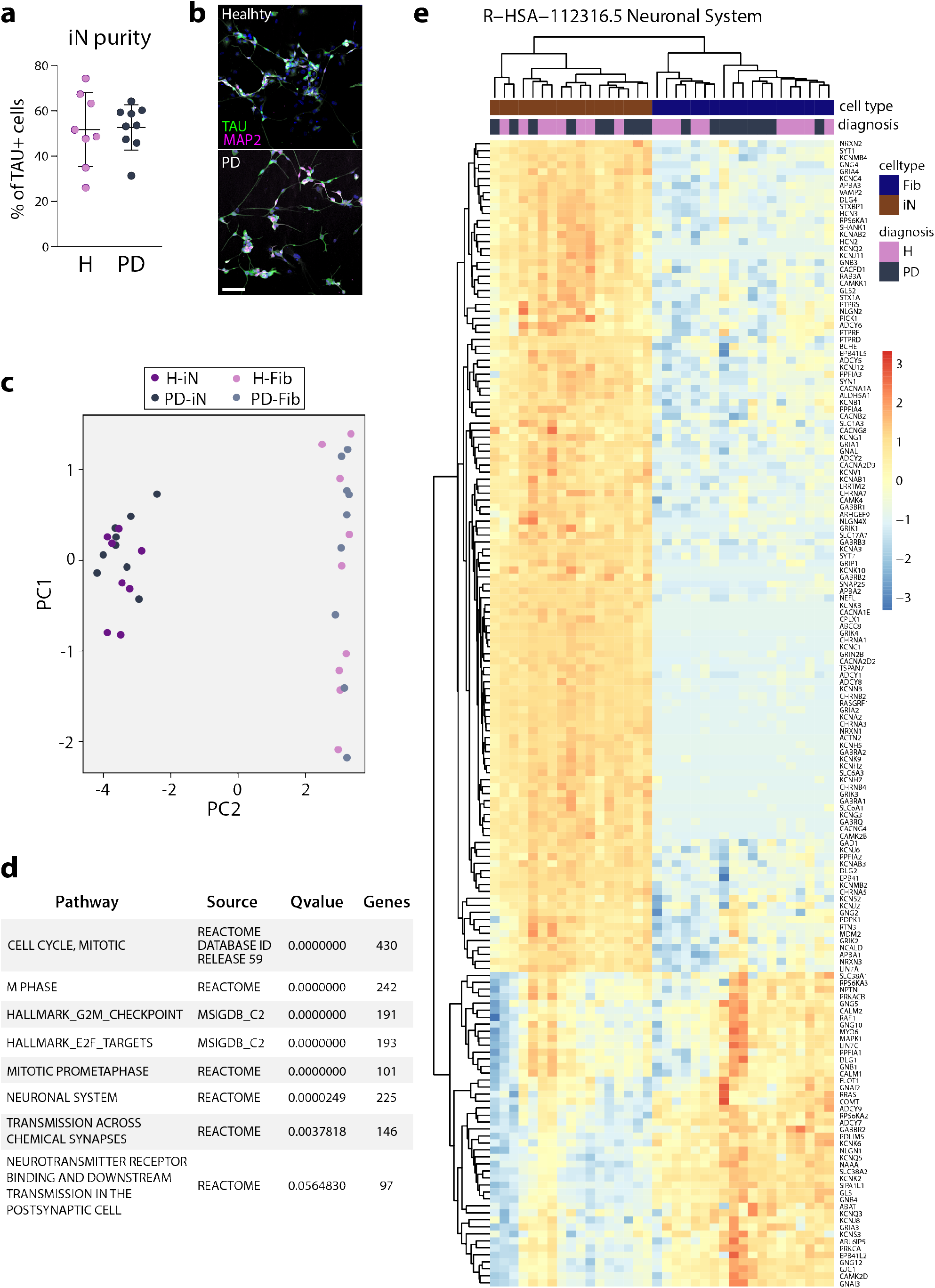
iN transcription profile. **a**, Quantification of the TAU-positive cell purity from a biological replicate of the cells sent for RNA-seq (mean average of 16,221 cells assessed per line).**b**, Double TAU-positive and MAP2-positive iNs from a biological replicate of the cells sent for RNA-seq. Scale bar = 100μm. **c**, Principal component analysis showing a separation of the reprogrammed iNs from the parental fibroblasts on PC2. **d**, Gene set enrichment analysis showing top ten most up and down regulated pathways in iNs compared to fibroblasts. **e**, Hierarchical clustering of RNA-seq samples, using Euclidean distance on normalized and log2-transformed read counts. Abbreviations: Fib: fibroblasts; H: healthy, iN: induced neurons; PD: Parkinson’s disease.

**Supplementary Fig. 8.**
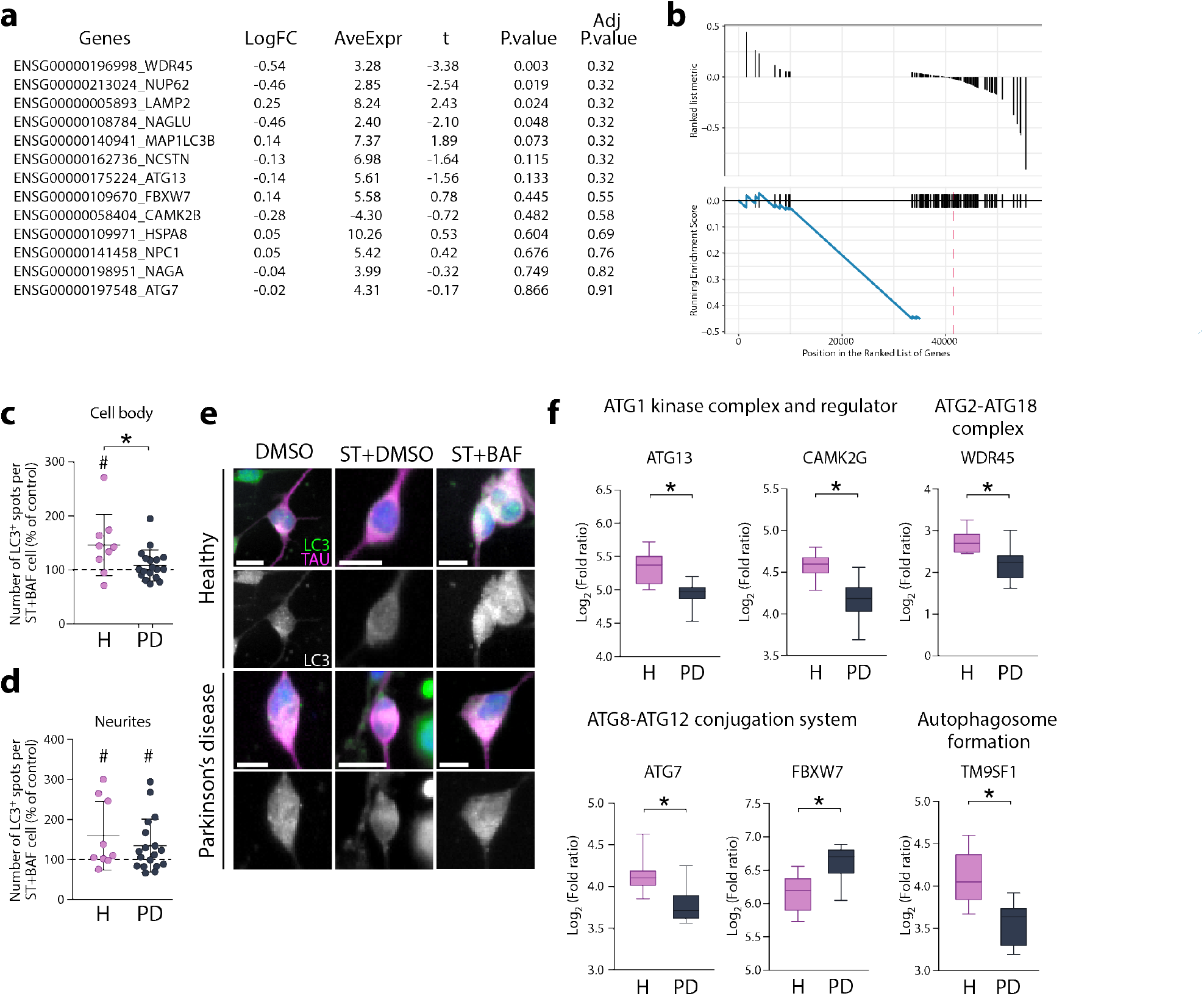
PD-iNs do not accumulate autophagosome structures following impairment of autophagy. **a,** LogFC and P. value table of autophagy and lysosome related genes in parental fibroblasts. Negative logFC means down-regulated in PD. Adj.P.Val” represents the P.value after transcriptome wide adjustment with the FDR method. (n=10 healthy and n=10 PD lines). **b,** GSEA plot of the “LYSOSOME” KEGG pathway in PD-iNs (Adj.P.Val = 0.028). **c**, Quantification of LC3-positive puncta in the cell body of TAU-positive iNs (mean average of 425 TAU-positive cells assessed per line, n=9 healthy and n=18 PD lines). Kolmogorov-Smirnov test: *P=0.0493, D=55.56. Two-tailed Wilcoxon matched pairs signed rank test, ^#^P=0.0371. **d**, Quantification of LC3-positive puncta in the neurites of TAU-positive iNs (mean average of 425 TAU-positive cells assessed per line, n=9 healthy and n=18 PD lines). H: two-tailed Wilcoxon matched pairs signed rank test, ^#^P=0.0371, PD: two-tailed paired t-test: ^#^P=0.0499, df=16. **e**, LC3-positive dot expression and spot detection analysis in TAU-positive iNs. Scale bar = 10μm. **f**, Boxplots of log2 fold changes in expression of genes associated with autophagy (n=10 healthy and n=10 PD lines) (adjusted P value < 0.05).

**Supplementary Fig. 9.**
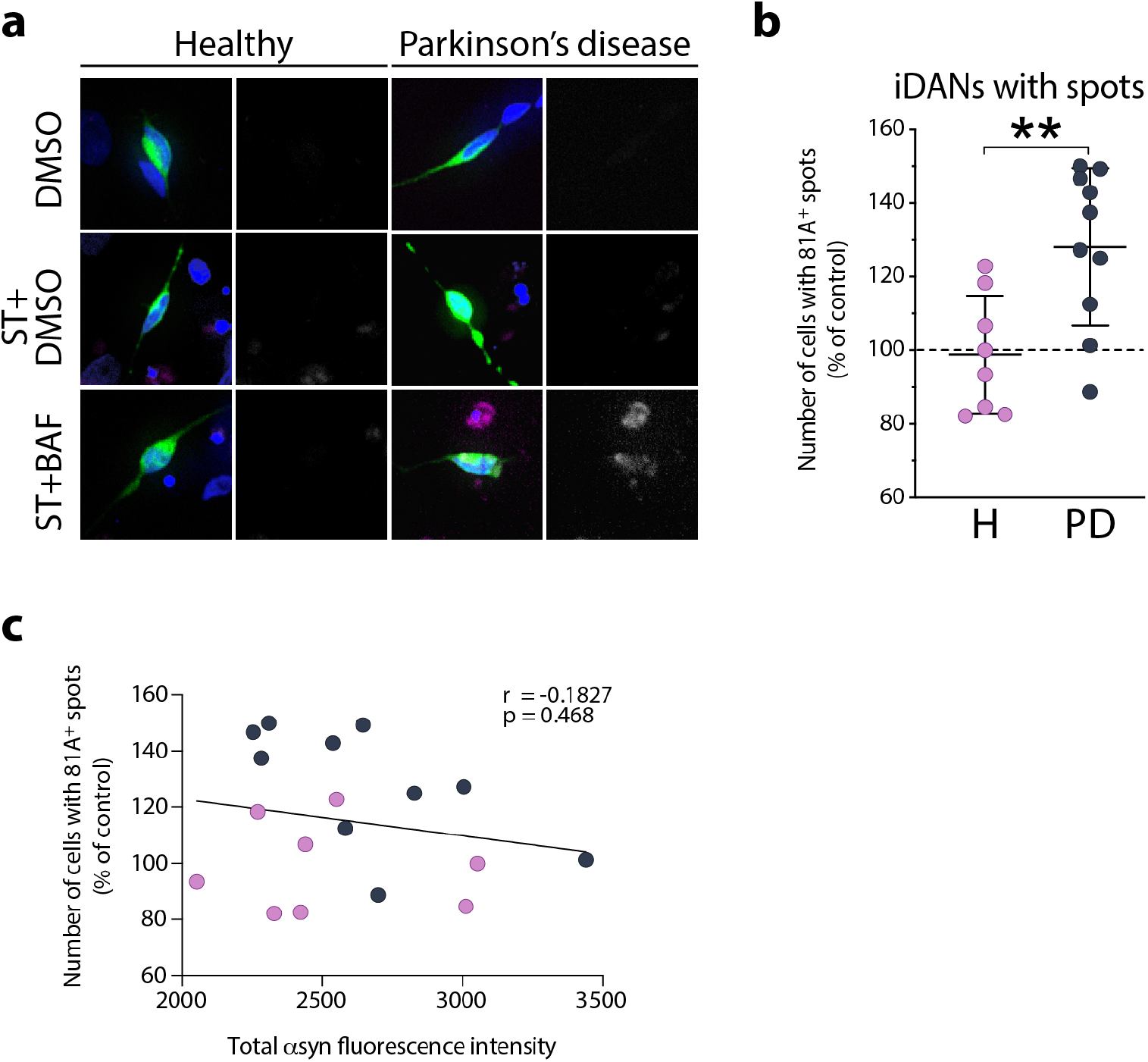
Autophagy impairments lead to an accumulation of phosphorylated αsyn in PD-iDANs. **a**, 81A-positive dot expression (in magenta) in TH-positive iDANs (green) directly reprogrammed from fibroblasts. **b**, Quantification of TH-positive iDANs with 81A-positive (pSer129 αsyn) puncta in the cell body (mean average of 93 TH-positive cells assessed per line, n=8 healthy and n=10 PD lines). Two-tailed unpaired t-test: **P=0.0054, df=16. Data were normalized as % of the control condition (% of starved+BAF/starved). **c**, No association between the number of 81A-positive puncta and the total asyn fluorescence intensity in iDANs (n=18 lines). Spearman’s rank correlation: P=0.468; 95% confidence interval: −0.6080 to 0.3242. Abbreviations: BAF: bafilomycin A1, H: healthy; PD: Parkinson’s disease, ST: starved.

**Supplementary Table 1.** List of Antibodies and primers

| **Antibody** | **Source** | **Species** | **Dilution** | **Catalogue number** | **Antibody registry number or PMIDs** |
| --- | --- | --- | --- | --- | --- |
| 81A | Gift from Kelvin Luk, University of Pennsylvania | Mouse | 1:10 000 | NA | 24931606,<br>24659240,<br>25732816 |
|  | Abcam | Mouse | 1:500 | Ab184674 | AB_2819037 |
| Alpha-synuclein | Abcam | Chicken | 1:1 000 | Ab190376 | AB_2747764 |
| Aldh1a1 | Abcam | Rabbit | 1:200 | Ab23375 | AB_2224009 |
| Collagen 1 | Abcam | Rabbit | 1:1 000 | Ab34710 | AB_731684 |
| GFAP | Millipore | Mouse | 1:1 000 | MAB3402 | AB_94844 |
| Anti-Histone H2A.X, phospho (Ser139) | Millipore | Mouse | 1:500 | 05-636 | AB_309864 |
| HSC70 | Abcam | Rat | ICC: 1:2 000<br>WB: 1:5 000 | Ab19136 | AB_444764 |
|  | Thermo Scientific | Mouse | 1:100 | MA3-014 | AB_325462 |
| LAMP2 | DSHB | Mouse | 1:100 | H4B4 | AB_528129 |
| LAMP2a | Abcam | Rabbit | ICC: 1:1 000<br>WB: 1:2 000 | Ab18528 | AB_775981 |
| LC3B | Sigma | Rabbit | ICC: 1:500<br>WB: 1:5 000 | L7543 | AB_796155 |
| MAP2 | Abcam | Chicken | 1:10 000 | Ab5392 | AB_2138153 |
| P62 | Abcam | Rabbit | ICC: 1:500<br>WB: 1:5 000 | Ab91526 | AB_2050336 |
| TAU HT7 | Thermo Fisher Scientific | Mouse | 1:500 | MN1000 | AB_2314654 |
| TAU | Agilent Technologies | Rabbit | 1:500 | A0024 | AB_10013724 |
| TH | Millipore | Sheep | 1:1 000 | Ab1542 | AB_90755 |
| VMAT2 | Sigma | Rabbit | 1:200 | AB1598P | AB_2285927 |

